# Genome-wide analysis of the cupin superfamily in cowpea (*Vigna unguiculata*)

**DOI:** 10.1101/2020.06.07.138958

**Authors:** Antônio J. Rocha, Mario Ramos de Oliveira Barsottini, Ana Luiza Sobral Paiva, José Hélio Costa, Thalles Barbosa Grangeiro

## Abstract

Cowpea [*Vigna unguiculata* (L.)Walp.] is an essential food crop that is cultivated in many important arid and semi-arid regions of the world. In this study the genome-wide database of cowpea genes was accessed in search of genomic sequences coding for globulins, specifically members of the cupin superfamily, a well-documented multigenic family belonging to the globulin protein class. A total of seventy-seven genes belonging to the cupin superfamily were found and divided into six families. We classify *V. unguiculata* genes into two subgroups: classical cupins with one cupin domain (fifty-nine proteins) and bicupins with two cupin domains (eighteen members). In addition, a search for cupin members in other closely related species of the fabaceae family [*V. angularis, V. radiatam* and *Phaseolus vulgaris* (common bean)] was performed. Based on those data, a detailed characterization and comparison of the cupin genes on these species was performed with the aim to better understand the connection and functions of cupin proteins from different, but related, plant species. This study was the first attempt to investigate the cupin superfamily in *V. unguiculata*, allowing the identification of six cupins families and better understand the structural features of those proteins, such as number of domains alternative splicing.

## 1. Introduction

Cowpea [*Vigna unguiculata* (L.)Walp.] is an important food crop that is cultivated in arid and semi-arid regions of Africa, Asia and Americas. In Brazil, it is mainly found in the northeast region, where it is a source of food for the population of that region (Ehlers *et al,* 1999).

Cowpea seed storage proteins are classified in four groups based on their solubility: albumins (water-soluble proteins), prolamins (alcohol-soluble), glutelins (acid or alkali-soluble) and globulins (diluted saline solution-soluble) (Osborne 1924). Globulins, in turn, are divided into two subgroups according to their sedimentation coefficients: 7S and 11S globulin-types, respectively known as vicilins and legumins (Ponzoni *et al,* 2018). Vicilins constitute the major source of nutrients during cowpea seed development (Kriz et al, 1999) and are composed of several isoforms encoded by multigenic families which are categorized based on the occurrence or not of enzymatic activity (Shotwell and Larkins 2012).

Furthermore, the cupin comprises a ubiquitious protein superfamily characterized by the presence of a conserved barrel domain (Dunwell, 1998). This domain has two conserved motifs of β-strands separated by a less conserved region composed by another two β-strands with an intervening variable loop (Dunwell *et al.*, 2000, 2001, 2002, 2003).

Cowpea 7S vicilins were found to contain two cupin_1 domains (bicupins), and β-vignins are the main representative of this protein class (Sales *et al.*, 1992, 2001). Cowpea β-vignins associate in trimers that form a carbohydrate-binding multiprotein. Each monomer possesses an oligosaccharide interacting site that confers specific carbohydrate-binding property to the oligomeric structure (Dunwell et al., 2002, 2003). These binding sites are located at the vertices of the triangle-shaped oligomer and the interaction between β-vignins and oligosaccharides, mainly through hydrogen bond interactions (a typical feature of carbohydrate-protein interaction), was suggested by computational simulations (Rocha *et al,* 2018).

In this study the genome-wide database of cowpea genes was accessed in search of genomic sequences coding for cupins, given that its represents a well documented multigenic family of globulins. A total of seventy-seven genes belonging to the cupin superfamily were found, which were then classified into six families by phylogenetic reconstruction methods. *V. unguiculata* cupin genes were categorized into two groups: classical cupins (fifty-nine proteins) and bicupins (eighteen members). In addition, a search for cupin members on other related species the fabaceae family (*V. angularis, V. radiatam* and *Phaseolus vulgaris*) was also performed. Based on these data, a detailed characterization and comparison of the cupin genes on the species analyzed was performed with the aim to better understand the connection and functions of cupin proteins from different, but related, plant sources.

## 2. Methods

### 2.1 Dataset

The *V*. *unguiculata* proteome IT97K-499-35 (genome assembly v1.0), available at the Phytozome database (http://phytozome.jgi.doe.gov/) (Goodstein et al., 2012), was accessed to search for proteins of the cupin superfamily. Furthermore, Rocha *et al,* 2018 were cloned six sequence denominate IT-81d-1053 (3R) and EPACE-10 3(S). 3 sequences resistance to *C. maculatus* and 3 susceptible to *C. maculatus,*

### 2.2 Sequences analysis

Analyses of the predicted the cupin superfamily proteins and identification of the cupin domain were performed using five different web servers. Pfam protein Database 2.0 (Finn et al, 2016), HMMER with Biosequence analysis using profile hidden Markov Models (Potter et al, 2018), SMART (Schultz et al, 2000) and Simple Modular Architecture Research Tool from the EMBL server and Conserved Domain tool from NCBI (CDD) (Marchler-Bauer., 2017). When applicable, only the result with the highest e-value was considered for analysis. BioEdit 7.2 software was used for edition (insertion and deletion) of amino acids sequences. The presence or absence of signal peptide was assessed with the SignalP 4,1 server (Petersen et al, 2011). The MEGA 7 (Tamakura *et al*, 2016) software was used for construction of phylogenetic tree using the Neighbor-Joining method (Saitou *et al,* 1987) with bootstrap values (1000 replicates).

#### 323 Protein structural model, docking and dynamic molecular

Proteins structural modeling, docking, and dynamic molecular were performed essentially as described elsewhere (Rocha et al., 2018).

## 3. Results and discussion

### 231- Cupin gene identification and analysis

We identified 77 gene sequences encoding proteins containing one or two copies of the cupin superfamily domain in the genome of *V. unguiculata* (Table S1). These sequences were grouped into six cupin families (cupin-1 to cupin-5 and cupin-8), and no sequences related to cupin-6 and cupin-7 families were found (Table S1).

The cupin-1 domain consists of a conserved barrel structure, and members of the cupin-1 family are represented by 11S and 7S seed storage globulins (termed legumins and vicilins, respectively) and germins (Dunwell et al, 1998). Legumins and vicilins are two-domain proteins (bicupins), whereas germins are single-domain molecules (monocupins) (Rocha et al, 2018) (Figure 1). β-vignins are the most abundant vicilins found in the genome of cowpea comprising 17 sequences (Table S 1 and 3), from which four are devoid of secretion signal peptides sequences: Vigun03g085800.1, Vigun03g085900.1, Vigun05g254700.1, vigun11g151800.1 (Table S2).

**Figure 1.**
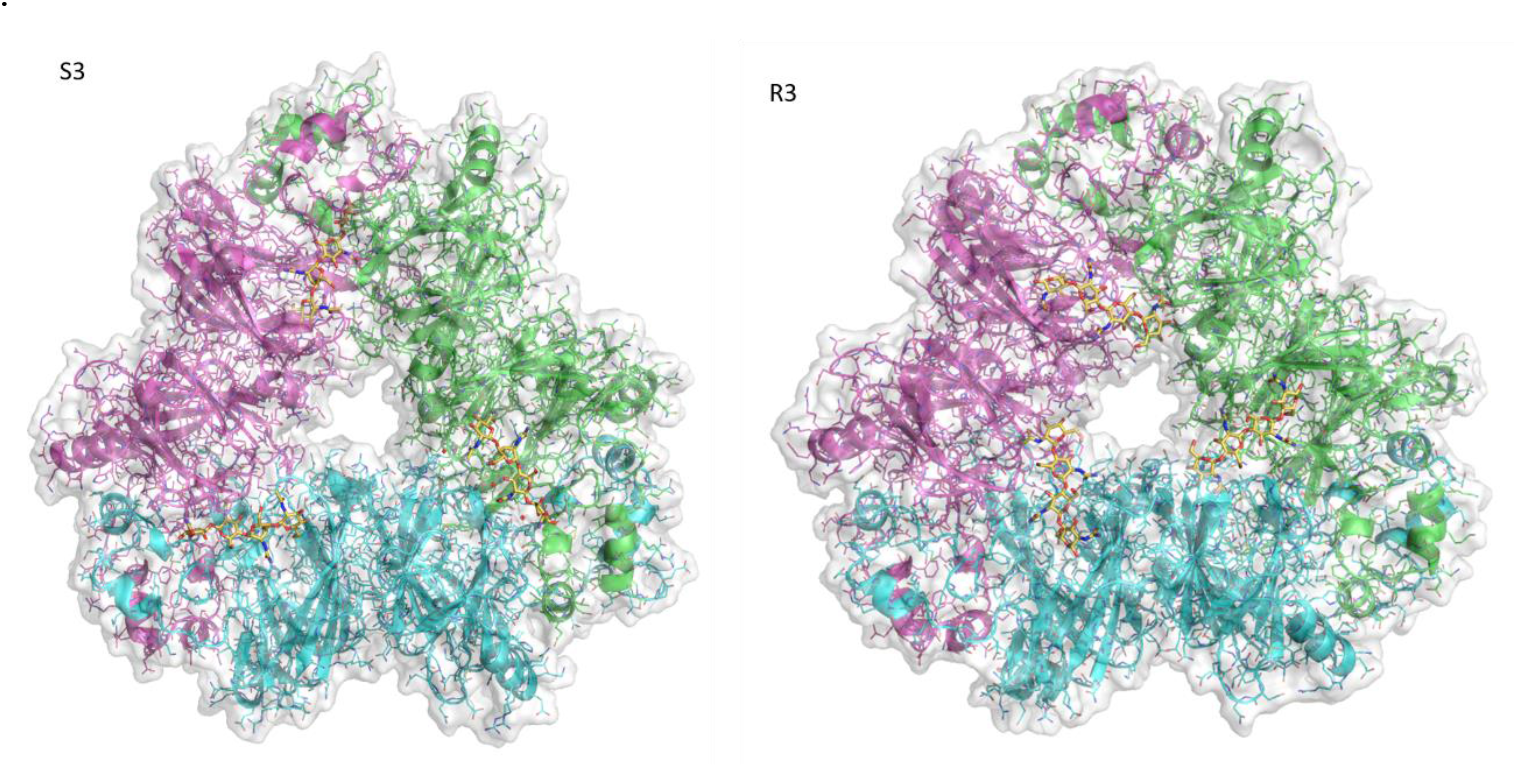
Ribbon diagrams of the β-vignin homotrimer models. Sequences Vicilin-S3-MG973243.1 (A) and Vicilin-R3-MG973246 (B) are shown. Subunits are colored pink, green and cyan. Chitooligosaccharide molecules [(GlcNAc) 4] docked in the chitin-binding sites of each oligomer are also shown as stick models (carbon, nitrogen and oxygen atoms are colored yellow, blue and red, respectively). For interpretation of the references to color, the reader is referred to the web version of this article

In a recent study based on computational simulations, Rocha et al (2018) demonstrated the presence of two cupin-1 domains in the primary structure of several β-vignin isoforms from two *V. unguiculata* genotypes (EPACE-10 and IT81D-1053) differing in the resistance to the bruchid beetle disease (*Callosobrcuhus maculatus*). In that study, the authors observed by computational simulations that β-vignin sequences presented a unique chitin-binding site (ChBS) in the N-terminal and in C-terminal ends (figure supp. 1). Those findings revealed the presence of three ChBS, which supports the hypothesis of the interaction of *V. unguiculata* β-vignins with the monosaccharide N-Acetyl-D-Glucosamine (GlcNac) and possibly its oligomeric derivatives, as observed for other bicupins in the present study (figure S1).

Cupin families 2 and 4 were represented by only one sequence with one cupin domain each. Cupin families 3, 5 and 8 were represented, respectively, by 8, 2 and 5 sequences with one cupin domain (Table S1 and 3).

We also identified sequences with 1 or 2 domains not belonging to the cupin-1 domain, such as auxin_BP, ARD, F-box-like, LacAB_rpiB. Some sequences contain one cupin domain and a second one not related to this superfamily. Sequences Vigun06g110700.1 (cupin-2 and LacAB_rpiB domain) and Vigun09g177900.1 (cupin-8 and F-box-like domains). Three sequences with domains Pirin and Pirin C were identified: Vigun03g399100.1, Vigun06g057200.1 and Vigun08g205900.1 (Table S1 and S3).

Other studies screening of the *Capsicum annuum* genome explored different proteins from the vicilin family, with attention to structural and functional features of Vic_capan sequences (vicilin of *C. annuum*) revealing that those vicilins belong to the cupin superfamily. Vic capan are known by their multifunctional enzymatic roles, ranging from epimerase and transferase in prokaryotes to oxalate oxidase and iron-binding nuclear protein (pirin) in eukaryotes (Dunwell, *et al* 2014). In the genome of pea (*Pisum sativum*), there are at least 18 genes encoding 7S vicilin divided into three small gene families, and 10 genes encoding legumins (Domoney and Casey 1985, Domoney et al. 1986). Furthermore, at least seven 11S globulin genes have been described in soybean (*Glycine max*), and they are arranged in three groups based on amino acid identity (Beilinson et al. 2002). More than 100 cupin sequences have been identified in few plant species, like *Oryza sativa*, *Vitis vinifera* and *Arabidopsis thaliana*. This finding highlights the extent of which cupins have been duplicated and diverged throughout the evolution in genomes of plants to carry out several functions. In addition, Sreedhar et al. (2016) purified a protease from rice (*Oryza sativa* L.), which was further denominated cupincin. This protein was included as a new member of the cupin superfamily.

The *V. unguiculata* cupins were also inspected for the presence of a secretion signal peptide. A total of 52 from the 77 sequences contain a predicted signal peptide sequence, as per the SignalP web server. Several of those sequences share conserved regions (Table S1 and S2).

In addition, the presence of alternative transcripts derived from alternative splicing in the mRNA with primary structure in the cowpea genome was also investigated. Four genes were found (Vigun03g085800, Vigun03g085900, Vigun05g251000 and Vigun07g160600), each one presenting 2 alternative transcripts. Two of these transcripts (Vigun03g085800 and Vigun03g085900) encodes each one polypeptides with identical amino acid sequences, whereas the transcripts of Vigun05g251000 and Vigun07g160600 encodes different proteins. The gene Vigun05g251000.1 encodes a protein with 377 amino acid residues, whereas Vigun05g251000.2 encodes a shorter version (341 amino acid residues) of the same protein, lacking the first 36 N-terminal residues in comparison to the larger isoform. The proteins encoded by Vigun07g160600 were identical, except in one isoform (Vigun07g160600.1; 226 amino acid residues), which was much longer than the other (Vigun07g160600.2; 222 amino acid residues), differing in 4 internal residues our insertion/deletion events.

In addition to *V. unguiculata* proteins, amino acid sequences belonging to other members of the cupin superfamily were also identified in closely related species: *Vigna angularis*, *Vigna radiata* and *Phaseolus vulgaris* (common bean) (Table S4 and S5

Eleven amino acid sequences of 7S vicilins, from which 10 are bicupins and one is a monocupin (ID: XP_017415489.1), were found in the genome of *V. angularis.* The same distribution pattern was found in *V. radiata* sequences, i.e. 10 sequences corresponding to bicupins and one monocupin, from a total of 11 sequences (Table S4). Regarding to 11S legumins, only 4 amino acid sequences were identified from the databank of cowpea, all sequences being of the bicupin type (Table S4).

Three web servers were used to investigate the *P. vulgaris* genome: NCBI CDD for search for conserved domains, SMART and HMMER (Table S5). Twenty sequences were found with the CDD web server. From these sequences, 12 were monocupins and 3 were bicupins. The remaining 5 sequences were not related to the cupin superfamily; instead, they were identified as proteins associated to the glutelin provisional domain. The SMART web server returned 20 sequences, all of which contain the vicilin-1 domain and 9 of those with the cupin-1 domain. These findings were similar to the HMMER web server results, which led to the identification of 20 sequences (10 bicupins and 10 monocupins) (Table S5).

As discussed before, the high copy number of globulin genes is not exclusive to leguminous plants, being also reported in non-leguminous ones. For example, in *Ficus pumila* (Moraceae family), six 11S globulin isoforms have been reported (Chua et al. 2008). Similarly, also in hemp (*Cannabis sativa* L.), a member of Cannabaceae family that possess seven 11S globulin (called edestin) genes, were identified and arranged into two groups (type1 and type2) based on differences on their primary structures (Docimo et al. 2014). The 11S edestin is the main storage protein representing approximatively 80 % of the total seed protein, whereas approximatively 13% of the water-soluble protein are 2S albumin.

### 3.2 Molecular Phylogeny

The phylogenetic classification of the cupin genes was performed initially only with sequences from *V. unguiculata* vicilins (Figure 2). Additionally, we analyzed sequences from 4 leguminous species from fabaceae family (*V. unguiculata, V. angularis, V. radiata* and *P. vulgaris*) (Figure 3). In the first phylogenetic tree, only 3 clades were identified between the *V. unguiculata* bicupins. The first clade includes 10 vicilin sequences obtained from cowpea database and 6 sequences of vicilins from cowpea with contrasting responses to the bruchid beetle (*Callosobrcuhus maculatus*) previously cloned by Rocha et al, 2018 (figure 2). In the second and third clades 4 and 5 vicilins are present, respectively, which correspond to the cupin_1 family. Others cupin families, such as cupin-2, 3, 4, 5, 7 and 8, are also present in this phylogenetic tree, as well as other sequences with non-cupin domain, such as Pirin-C, ARD and Axin_BP.

**Figure 2.**
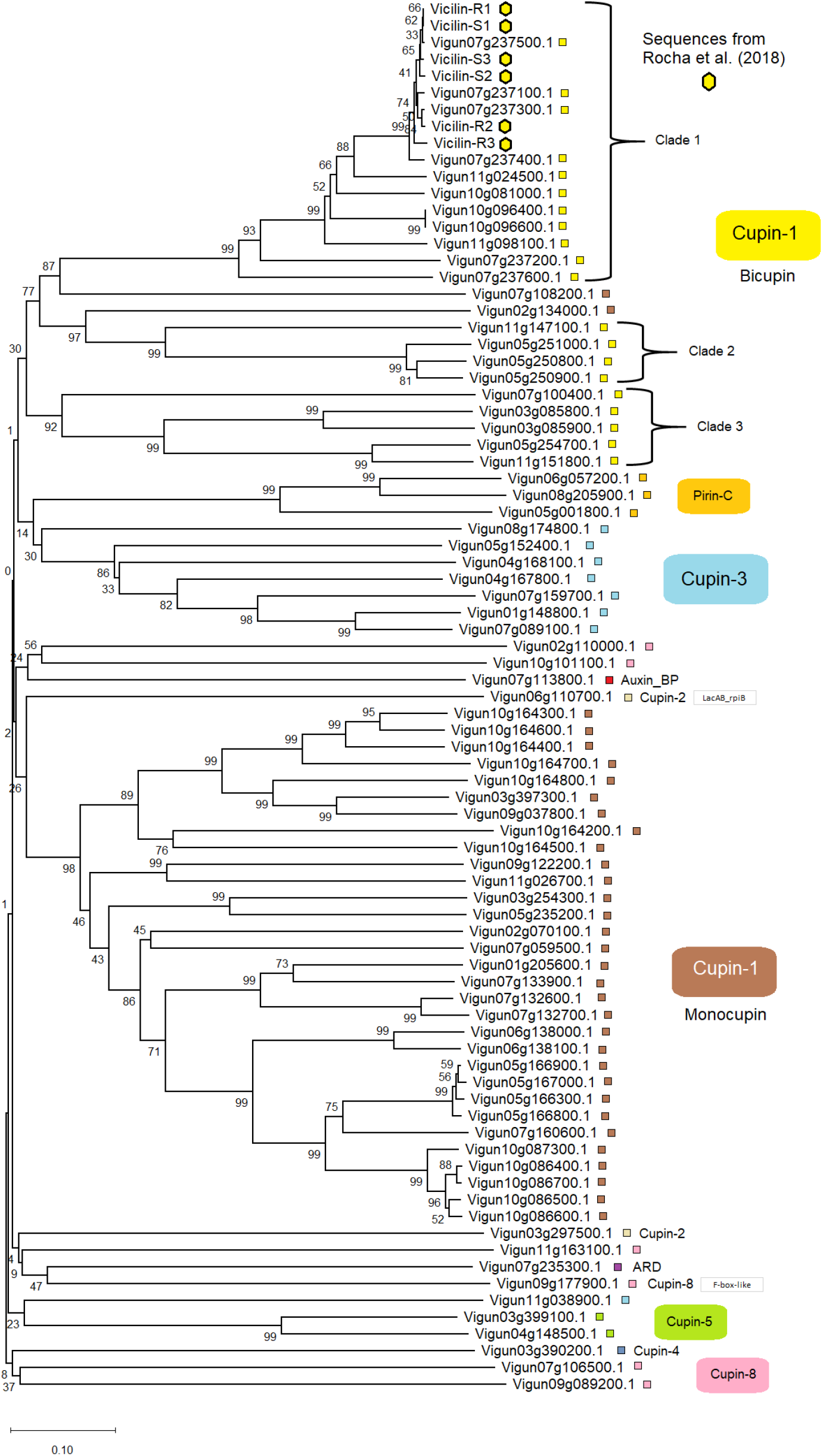
Molecular phylogeny of *V. unguiculata* cupins. The different cupin families are highlighted in several colors and include monocupin and bicipuns (vicilins).

**Figure 3.**
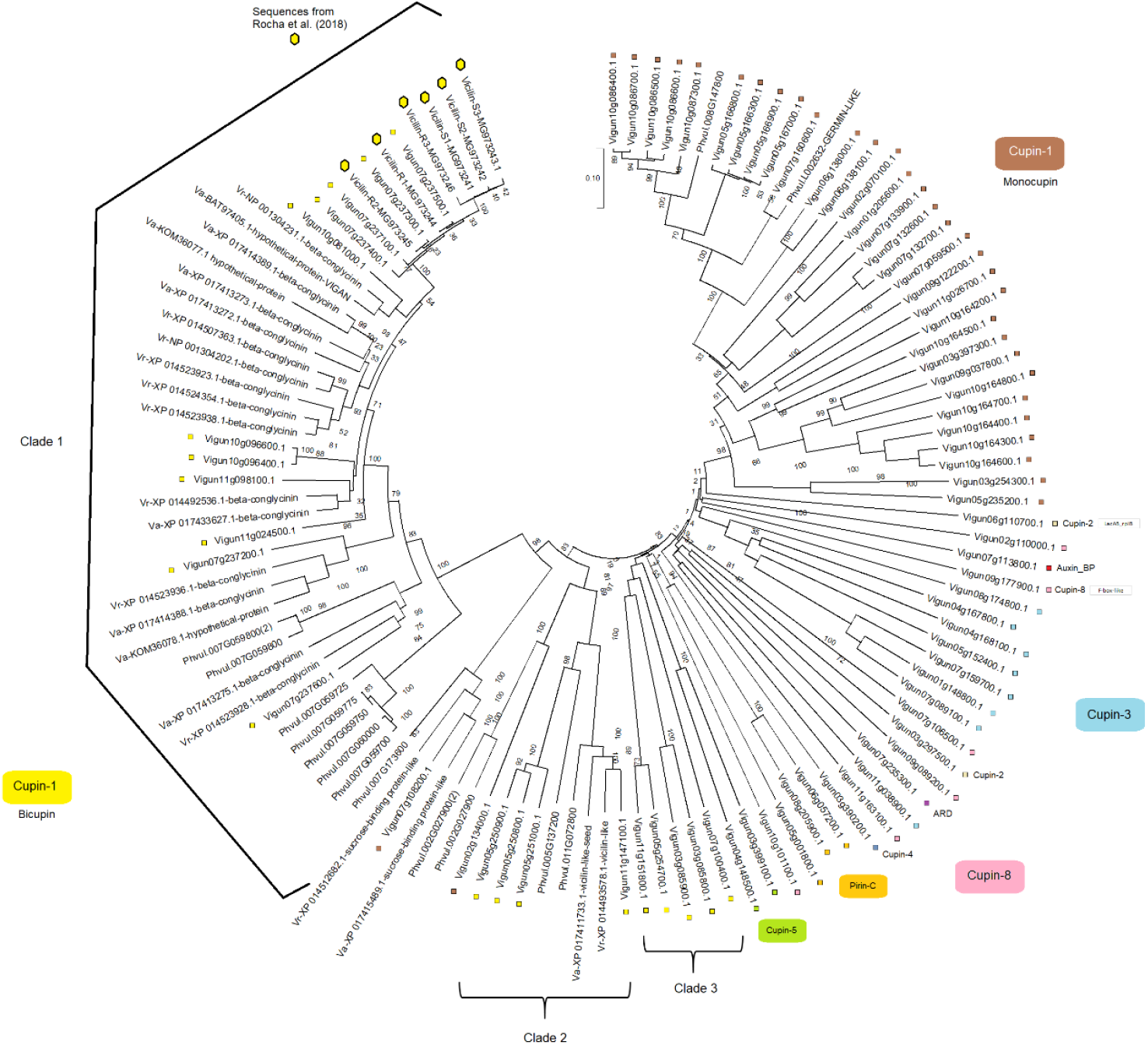
Molecular phylogeny of *V. unguiculata, V. angulares, V. radiata and P. vulgaris* cupins. This tree depicts clade 1 with *V. unguiculata* bicupins from the cupin-1 family, as well clades 2 and 3 that were also identified. The largest group in terms of *V. unguiculata* sequences is the one formed with cupin-1 monocupins.

With respect to the analysis including the 4 selected plant species, the results showed that 3 clades were classified in accordance with position of the 19 amino acids sequences of bicupins of *V. unguiculata* (Figure 3). Again, clade 1 contains 9 bicupins obtained from the *V. unguiculata* database, as well as the 6 sequences cloned by Rocha et al (2018). Clade 2 contains 5 sequences from *V. unguiculata* bicupins. Clade 3 contains 4 *V. unguiculata* bicupins and other bicupins from *V. angulares, V. radiata* and *P. vulgaris* (Figure 3).

All other amino acid sequences from *V. angularis, V. radiata* and *Phaseolus vulgaris* are bicupins that were not grouped with *V. unguiculata* bicupins. Moreover, the 34 *V. unguiculata* monocupins identified (non-vicilins) form one larges clade. Three pirin domain-containing sequences (pirin and pirin C), which belong to the cupin-2 family were also grouped in this analysis. Other research groups have studied human pirin sequences and have obtained their crystal structure, which is an iron-binding nuclear protein and transcription cofactor (Pang *et al*. (2004). In accordance with the first phylogenetic tree (Figure 1), in the second tree (Figura 3) also grouped cupin sequences belonging to the same family (1, 2, 3, 4, 5, 7 and 8), as well as sequences with no cupin domain, such as Pirin-C, ARD and Axin_BP.

### Conclusion

In this study we identified six cupin families in the *V. unguiculata* genome, followed by the analysis of copy number of 7S vicilins and/or 11S legumin sequences from *V. unguiculata* and other leguminous plants, such as *V. angularis*, *V. radiata, P. Vulgaris*. This led to the identification of numerous cupin sequences in each species. Furthermore, the kind of and number of domains that are present in each sequences are described; and genes that are originated due the alternative splices are proposed. In summary, our results represent the first attempt at a thorough characterization of cupin sequences in important leguminous plants. This will support future studies to elucidate their biological role, which are still needed for a complete understanding of the cupin superfamily function, including *V. unguiculata* β-viginins with two cupin-1 domains that are the major constituent of cowpea vicilins.

## Acknowledgements

This work was supported by grants from “Conselho Nacional de Desenvolvimento Científico e Tecnológico” (CNPq) and “Coordenação de Aperfeiçoamento de Pessoal de Nível Superior” (CAPES). AJR was recipient of Doctoral Fellowships from CAPES and CNPq. AJR, ALSP and JEMJ wrote the manuscript. JHC done the phylogenetic tree.

## SUPPLEMENTARY MATERIAL

**Table S1.**
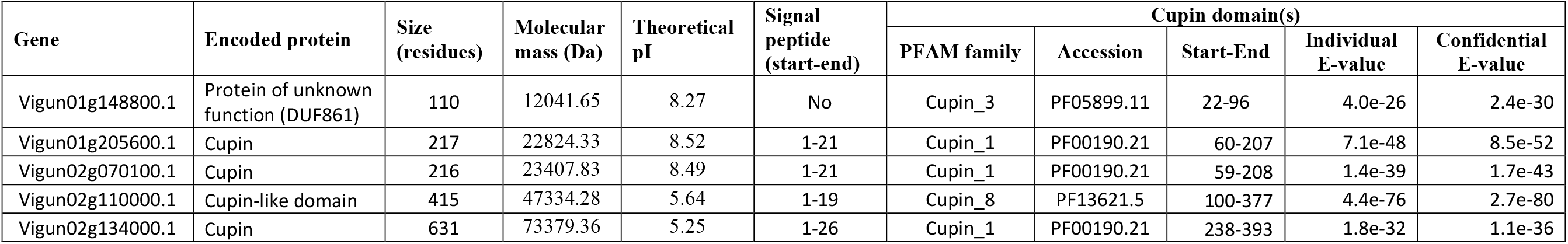
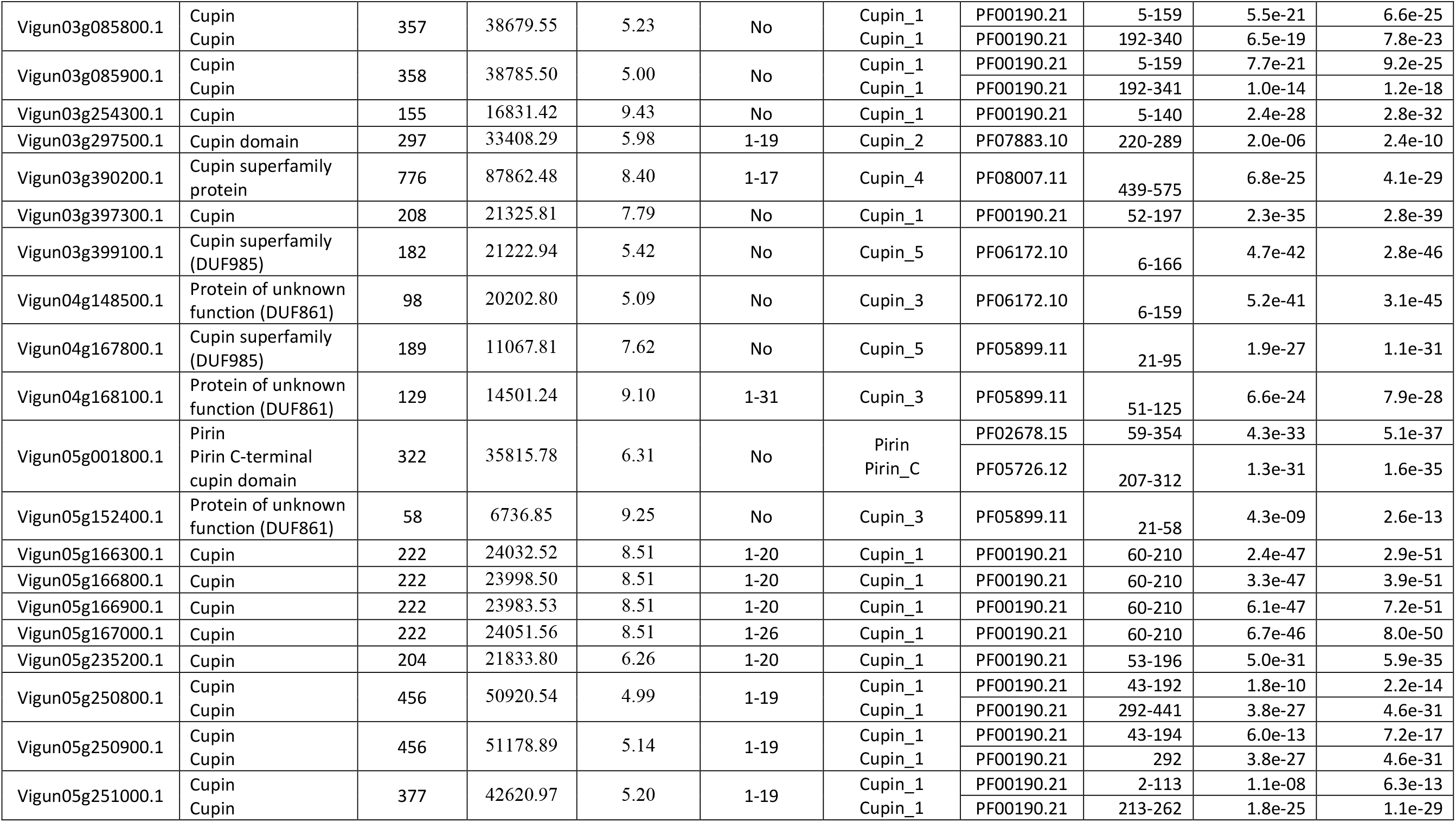
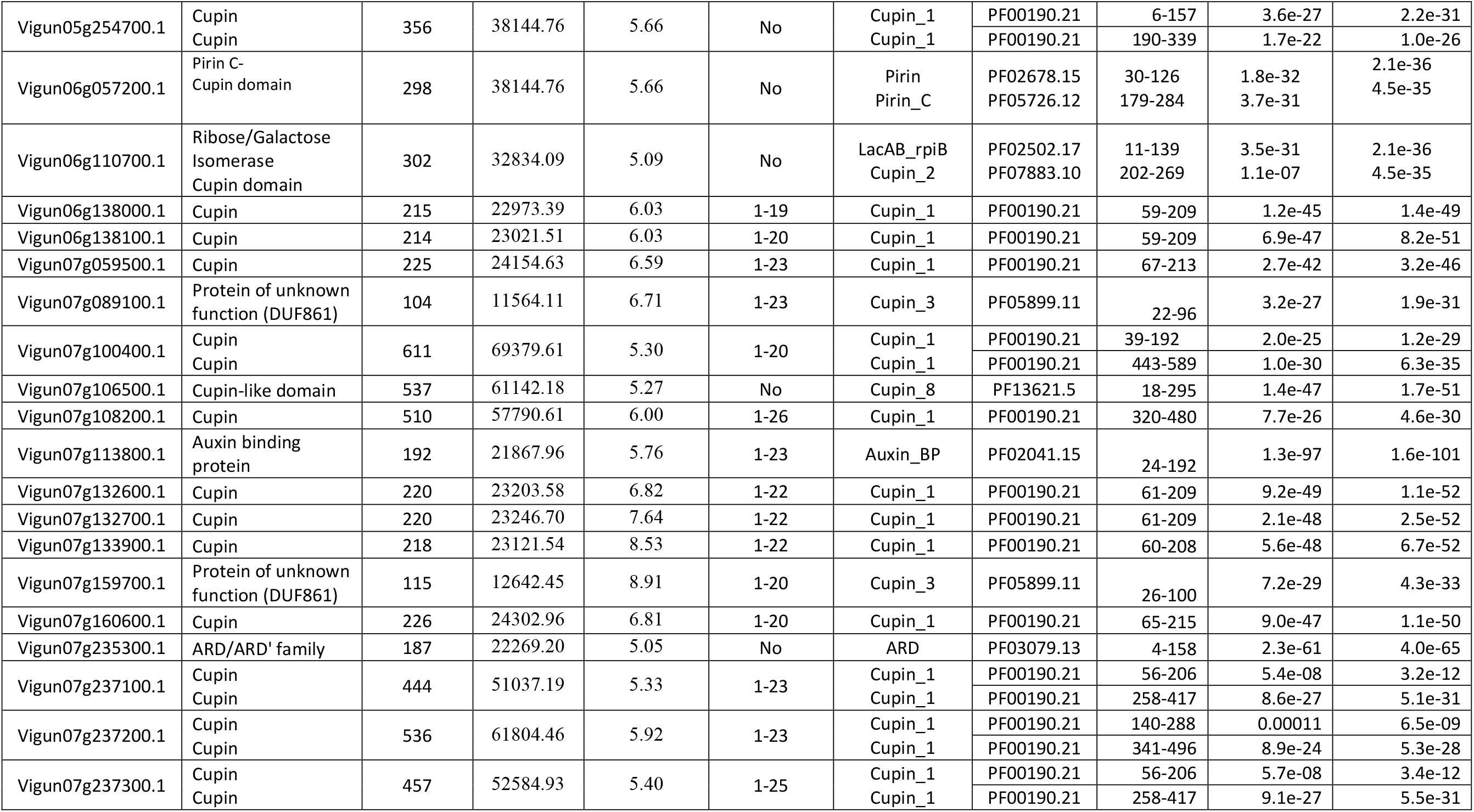
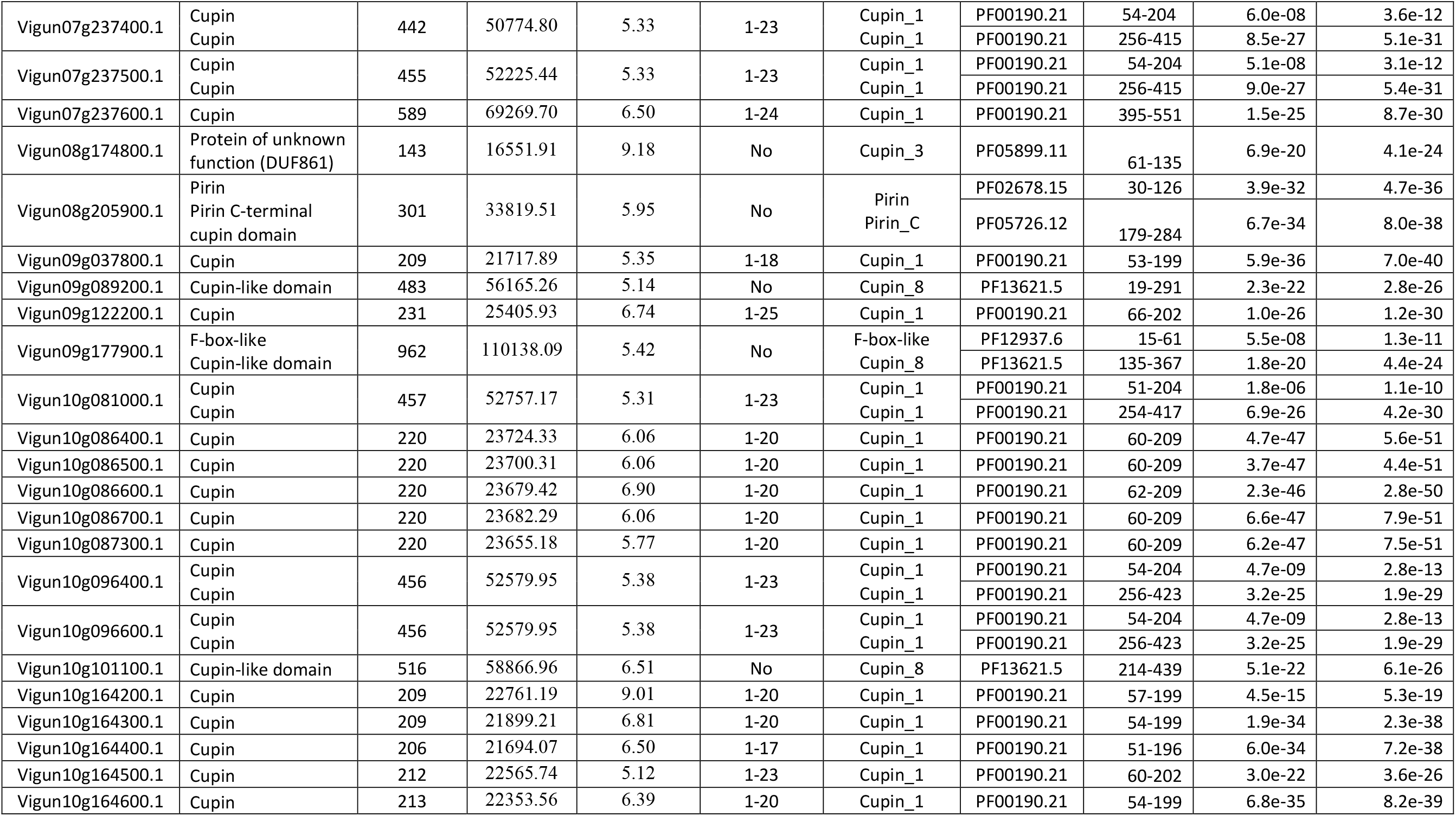
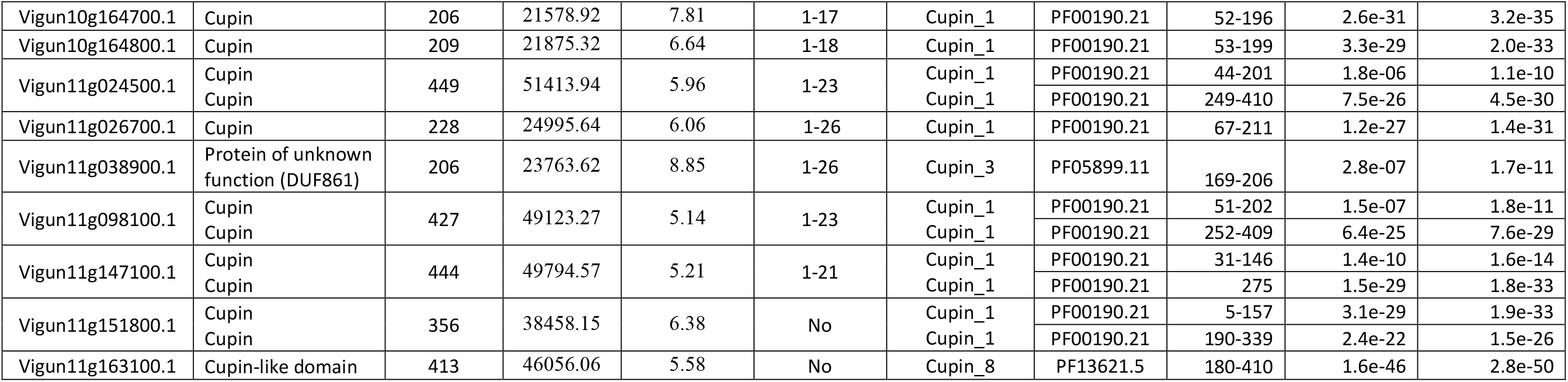
Cupin superfamily proteins from cowpea (*Vigna unguiculata*)

**Fig. S1.**
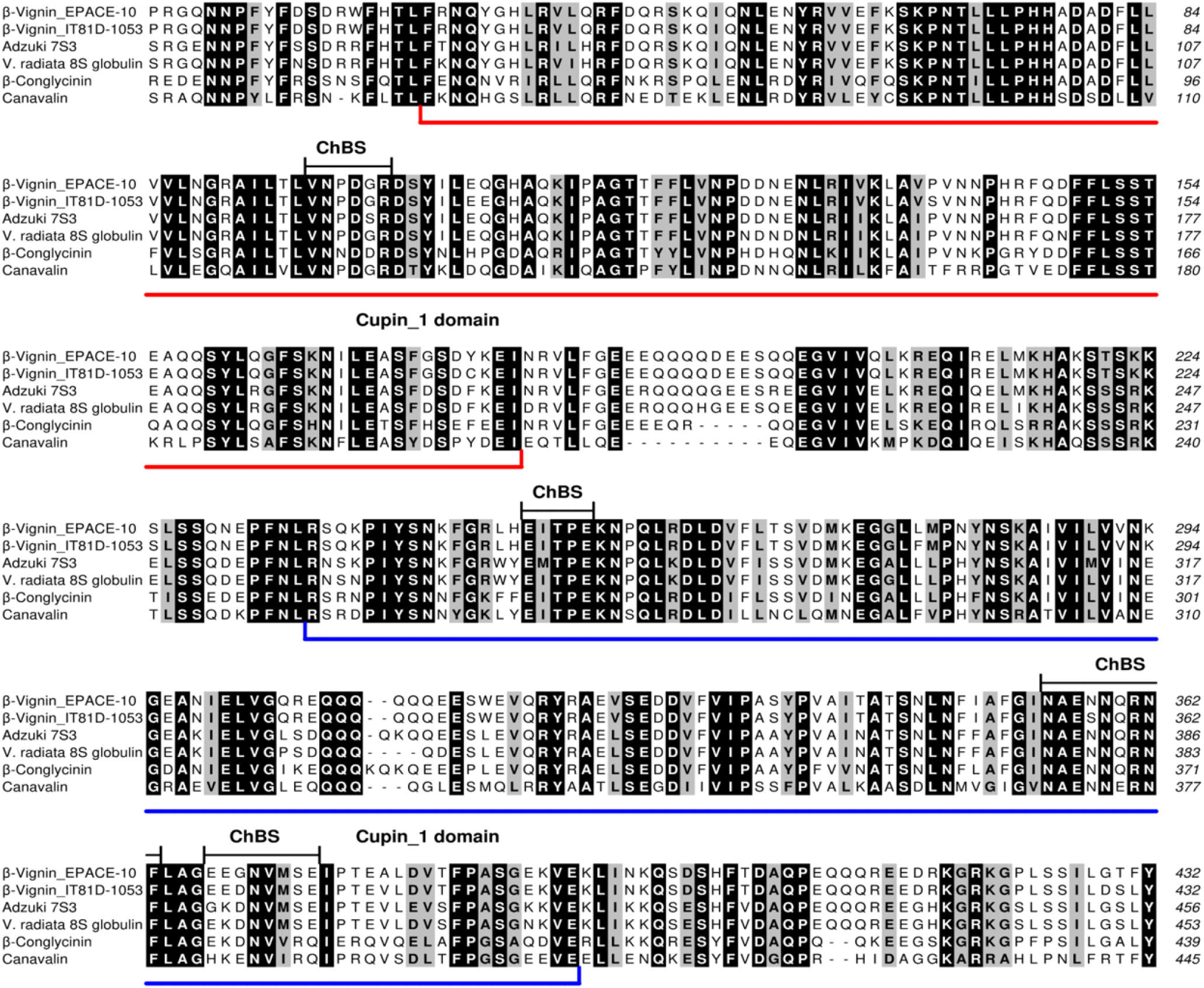
Multiple sequence alignment of the amino acid sequences of β-vignin with the primary structures of representative vicilin-like 7S globulins. Amino acid sequences of β-vignin obtained from V. unguiculata genotypes EPACE-10 (sequence S2) and IT81D-1053 (sequence R2) were aligned with those of V. angularis (adzuki bean) 7S globulin-3 (Adzuki 7S3;UniProtKB accession number: A0A0S3SX36), V. radiata 8S globulin (UniProtKB accession number: Q198W3), β-conglycinin (from Glycine max; UniProtKB accession number: P25974) and canavalin (from Canavalia ensiformis; UniProtKB accession number: P50477). Segments in the primary structures of β-vignin that were shown to contribute to their chitin-binding site (ChBS), as evidenced by computational simulations, are indicated. The alignment was edited using the program ALINE.

**Table S2.**
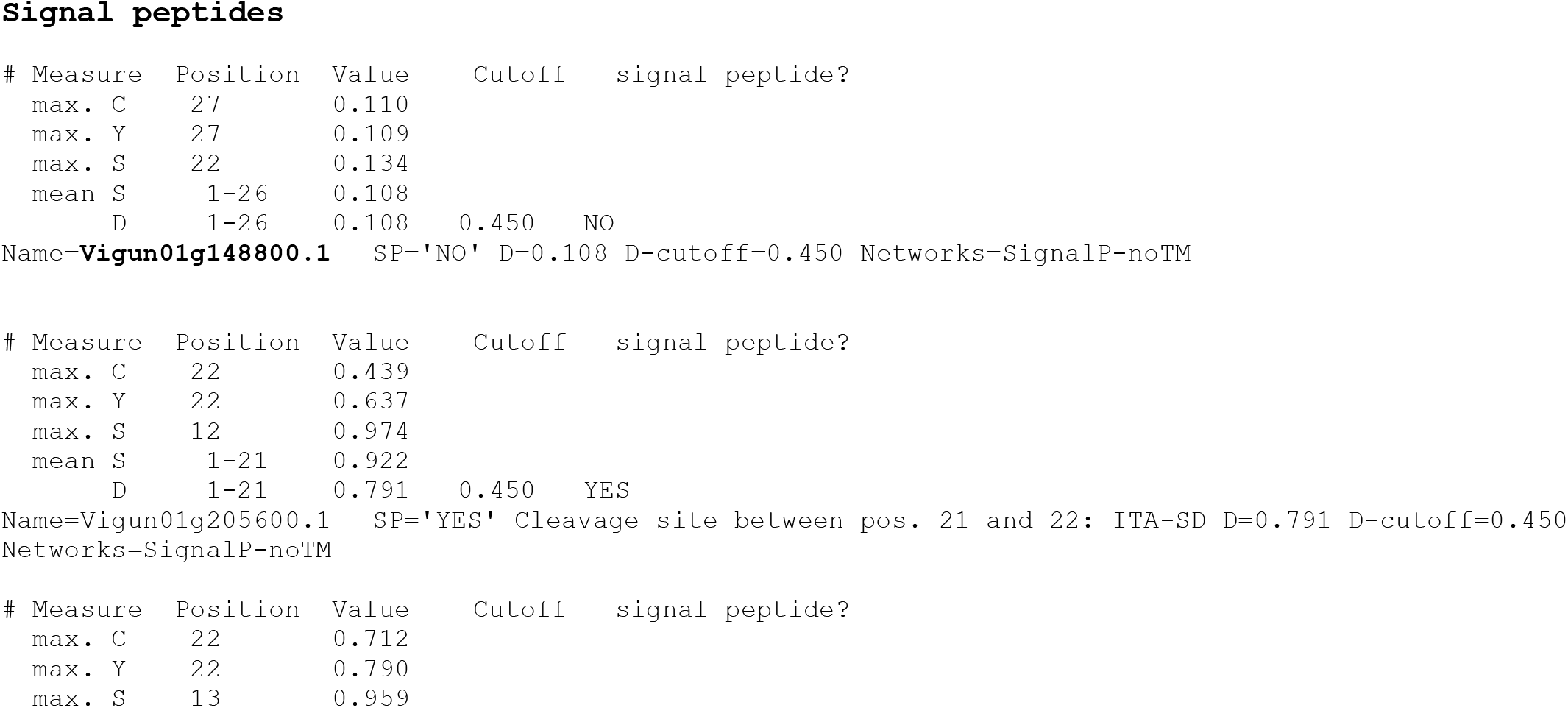
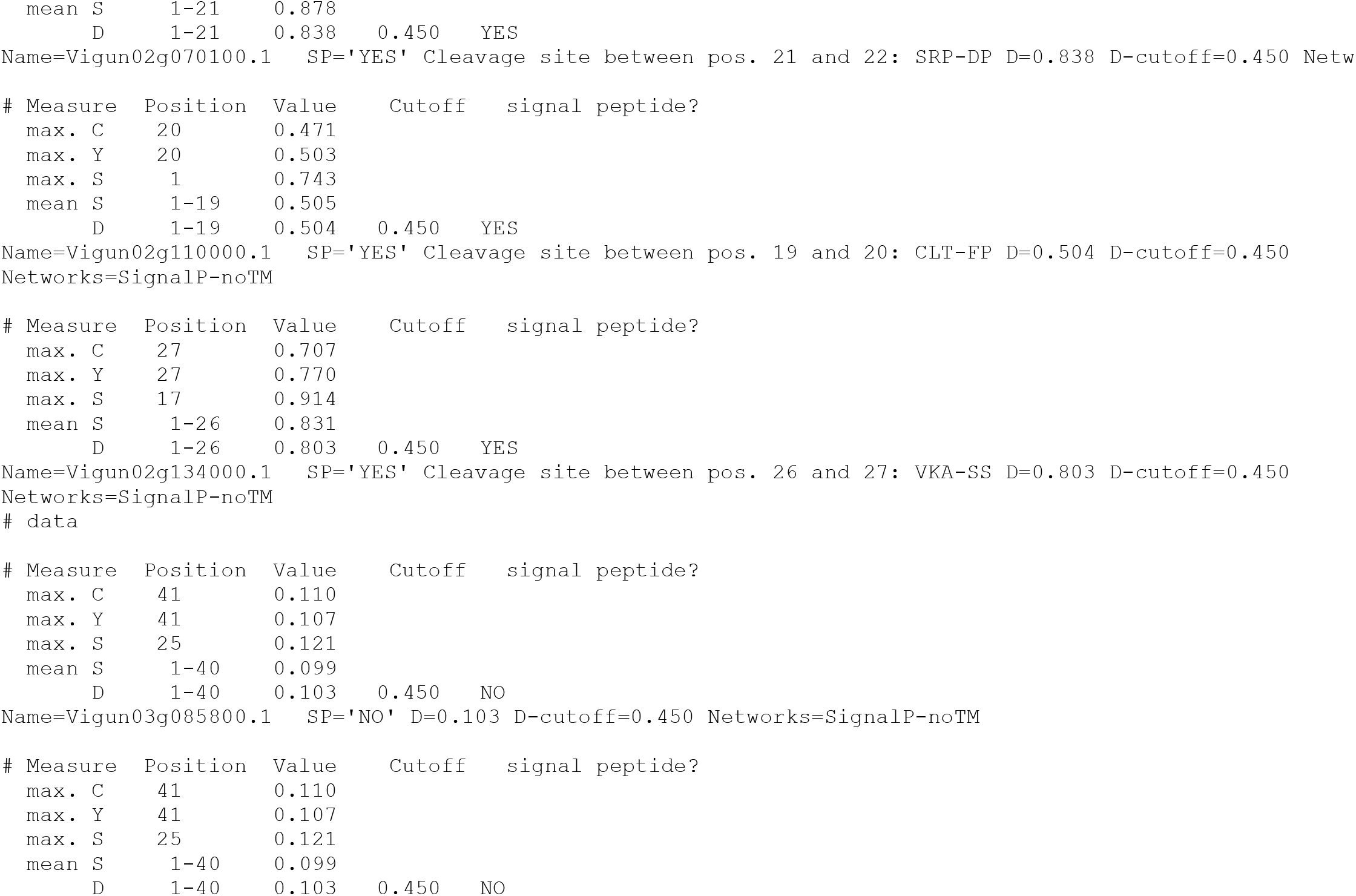
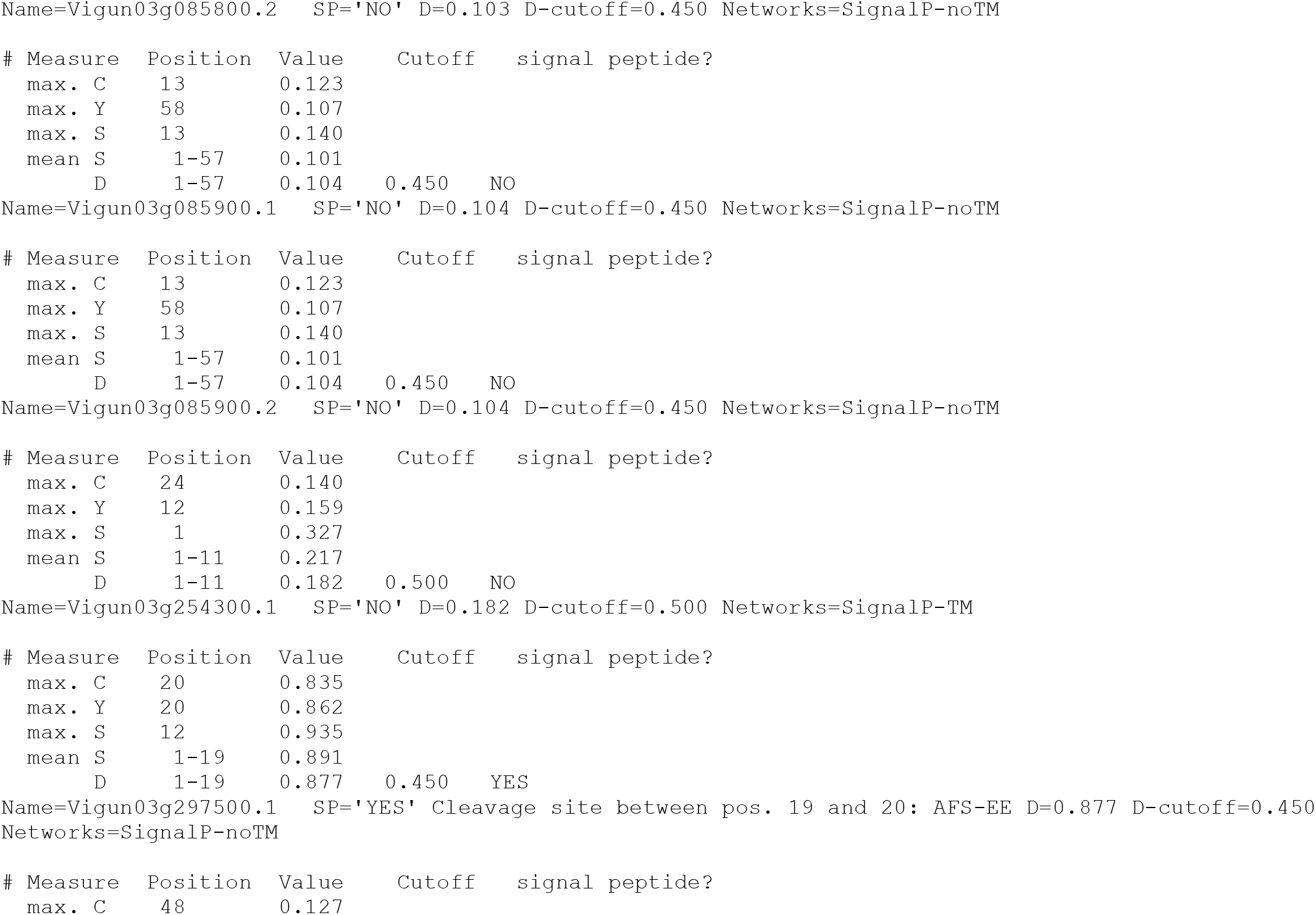
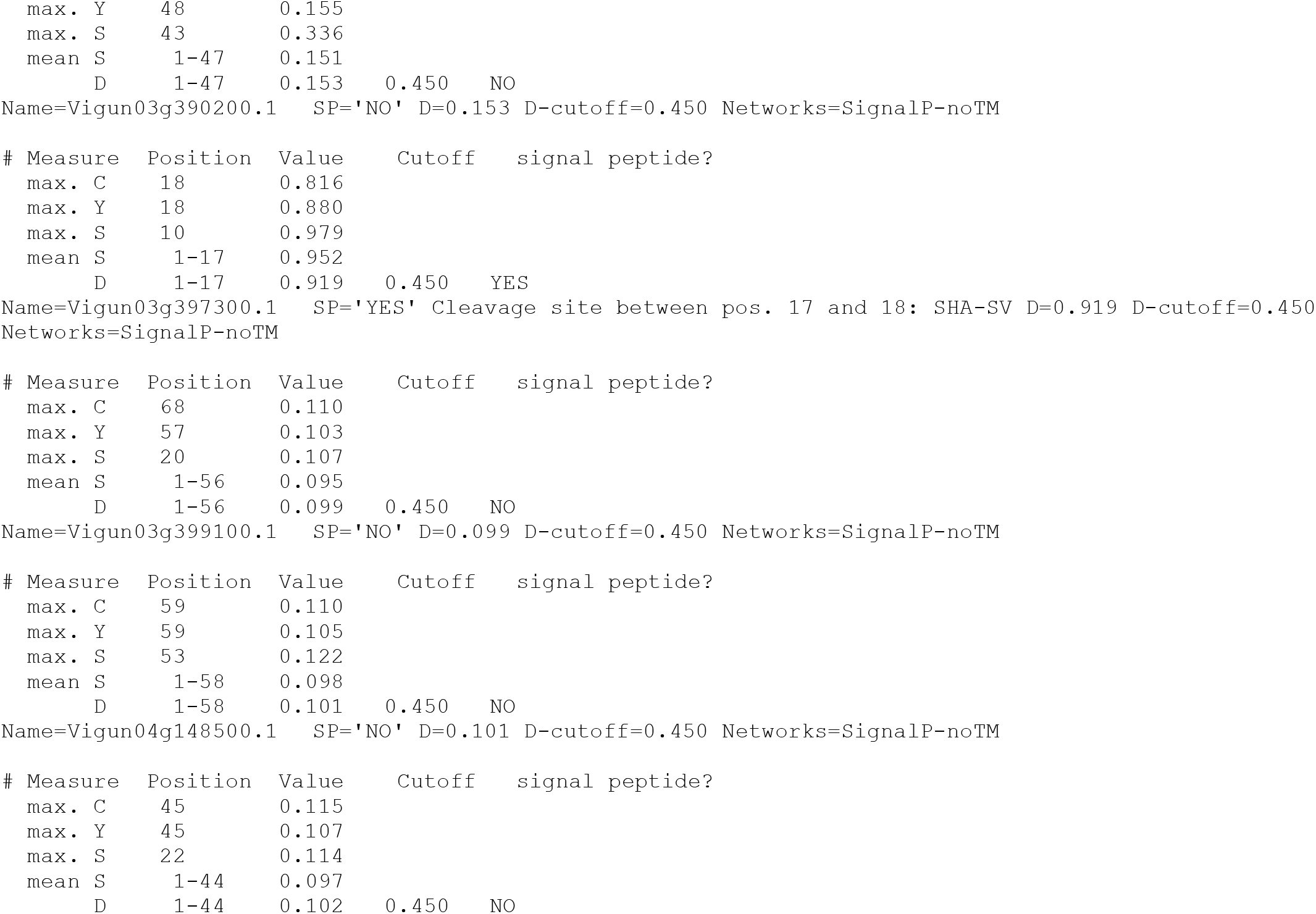
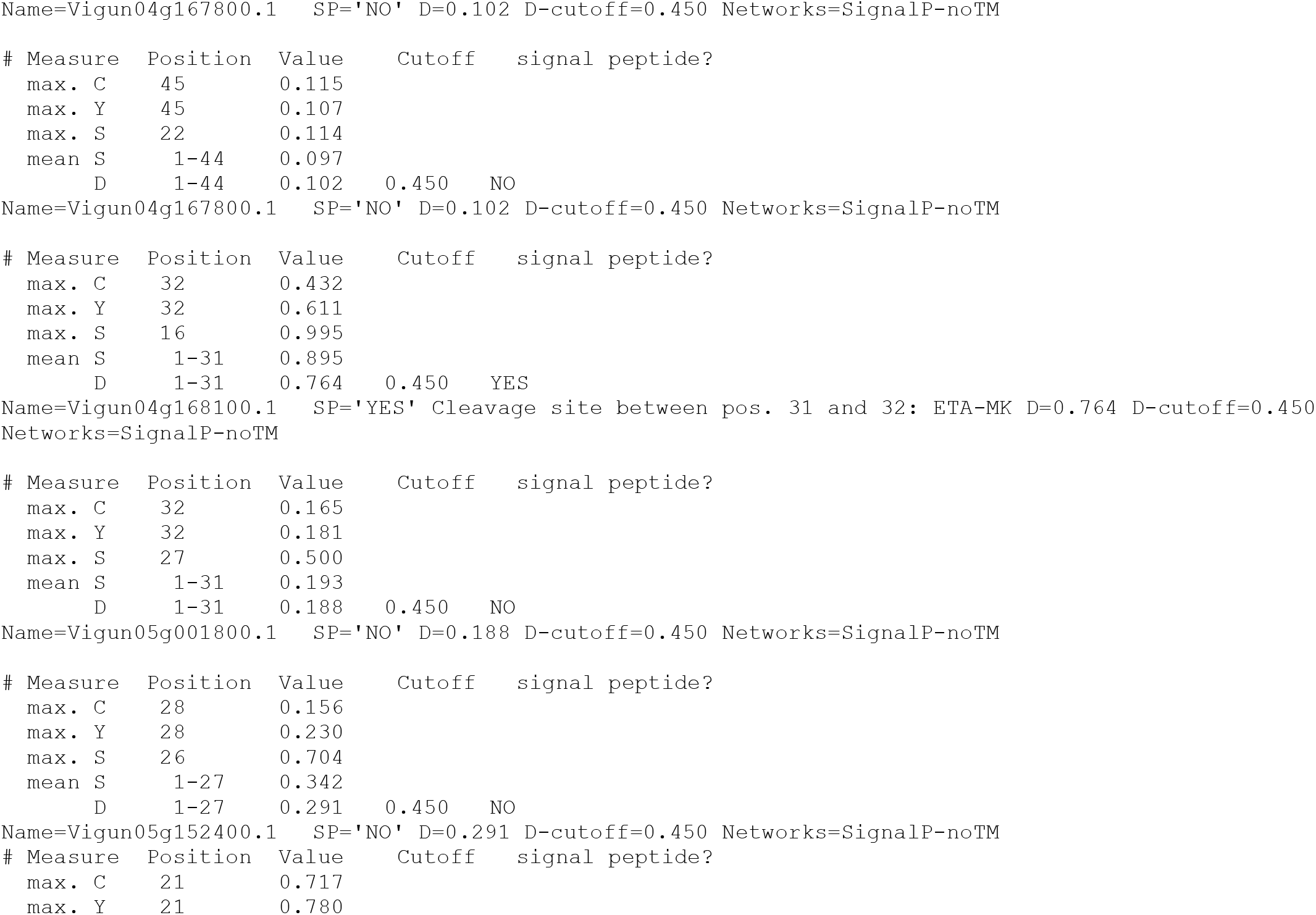
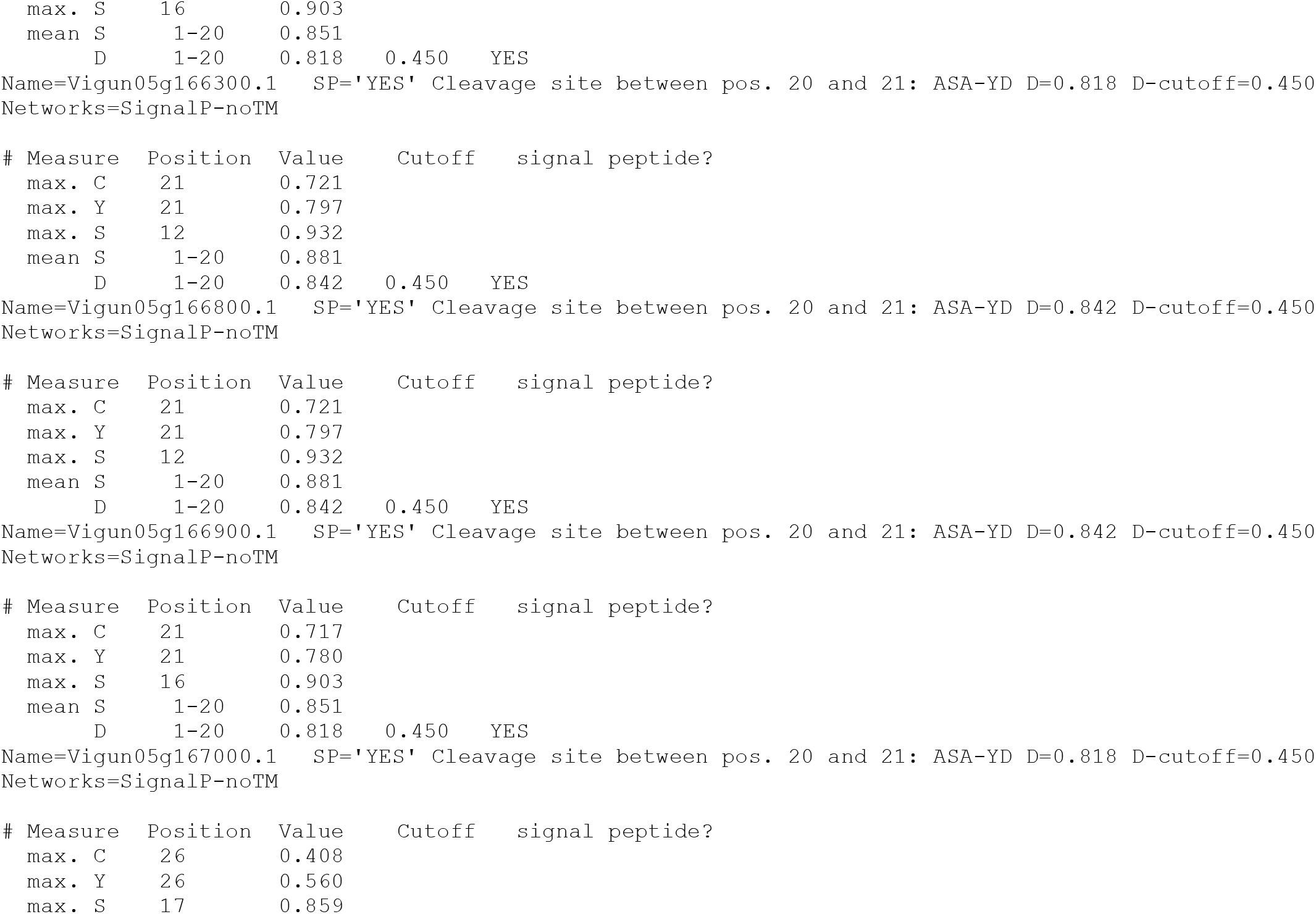
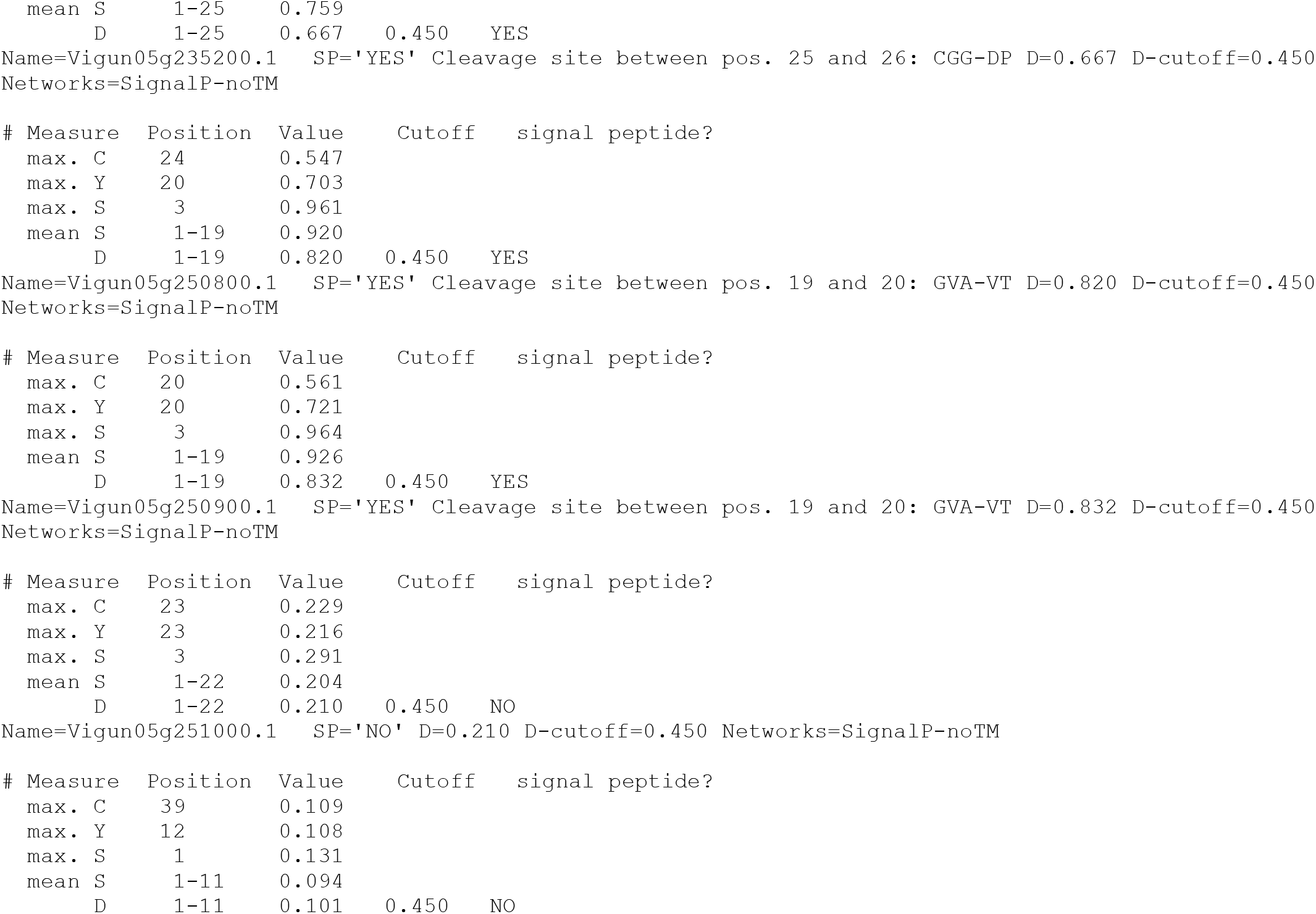
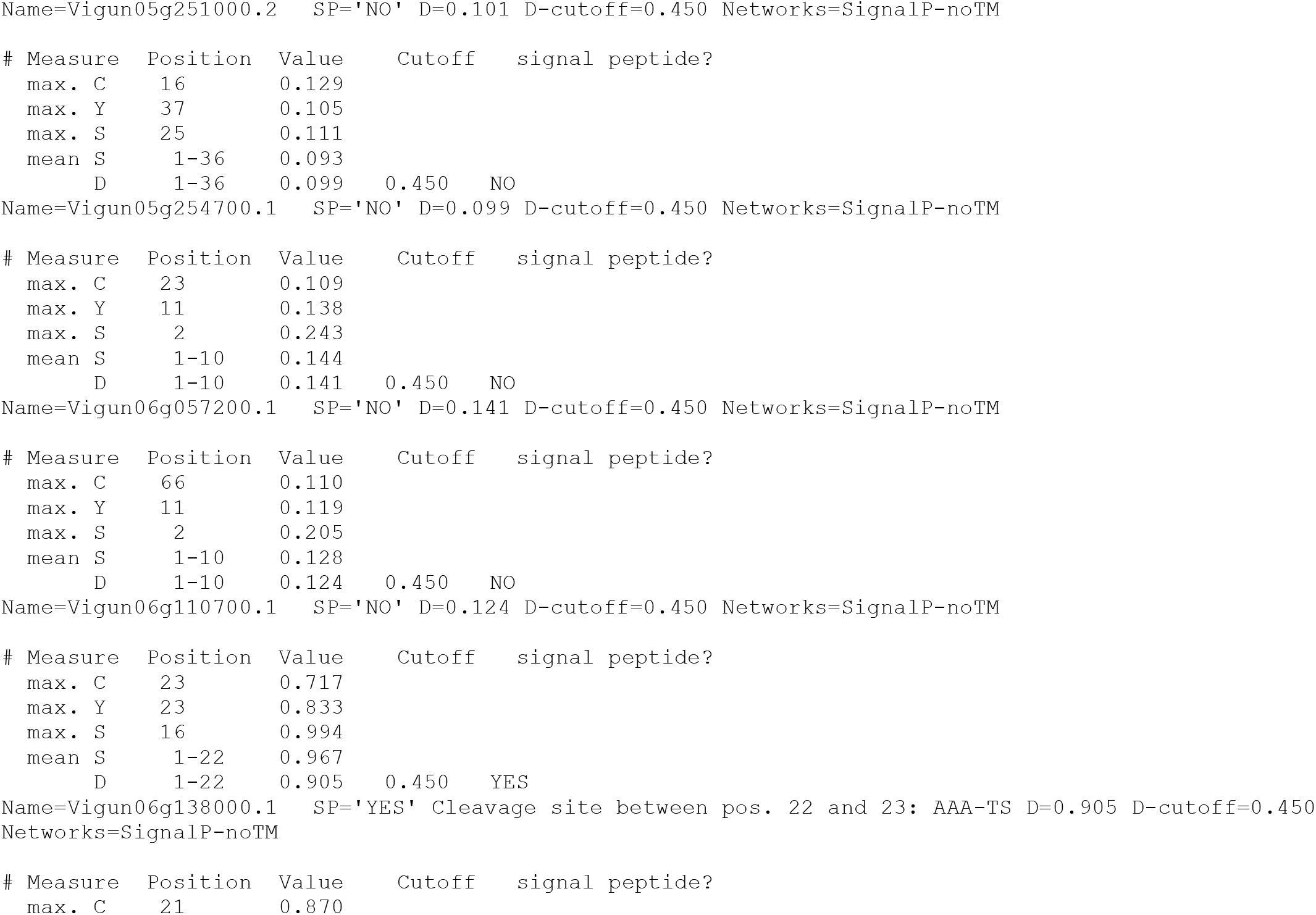
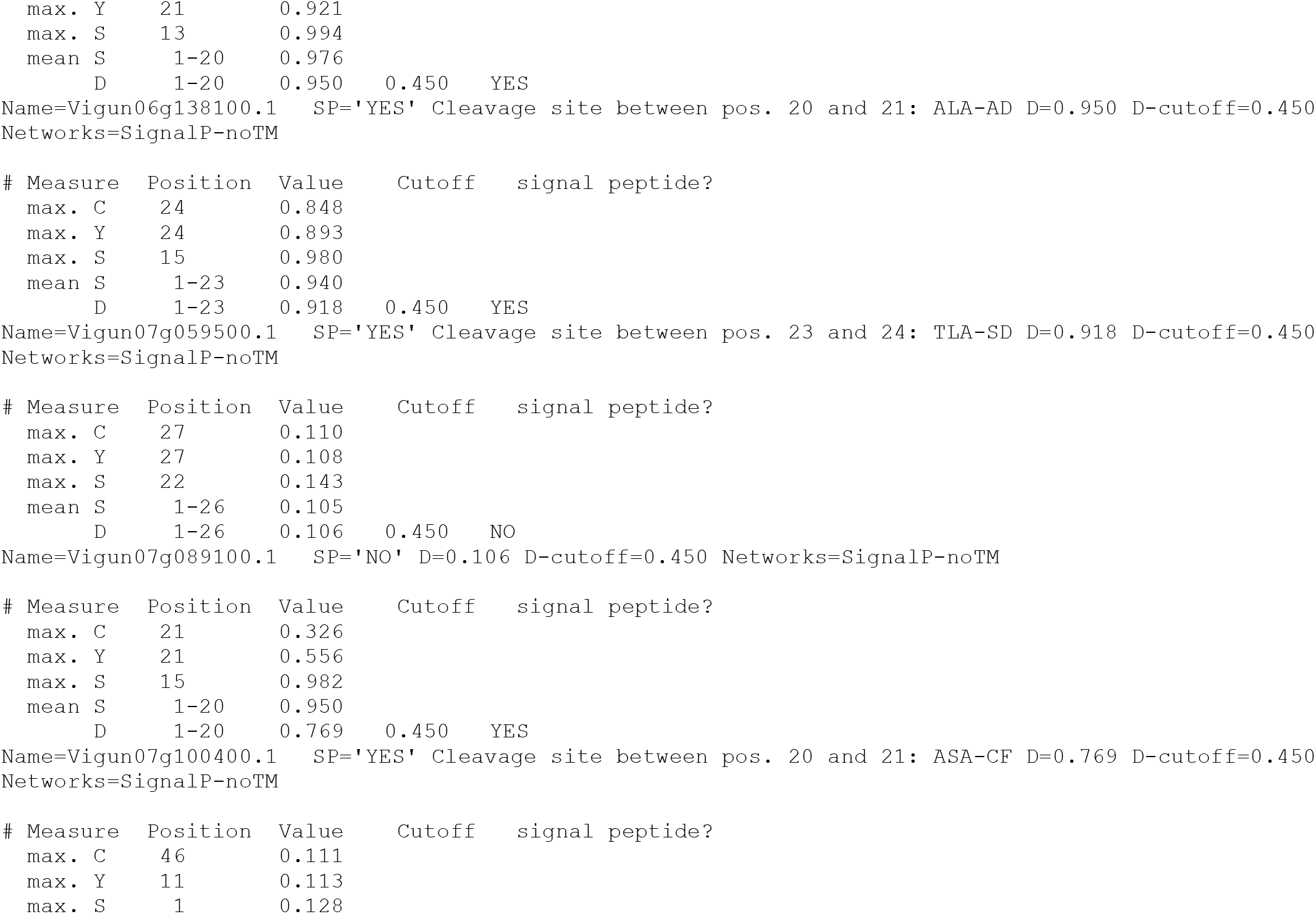
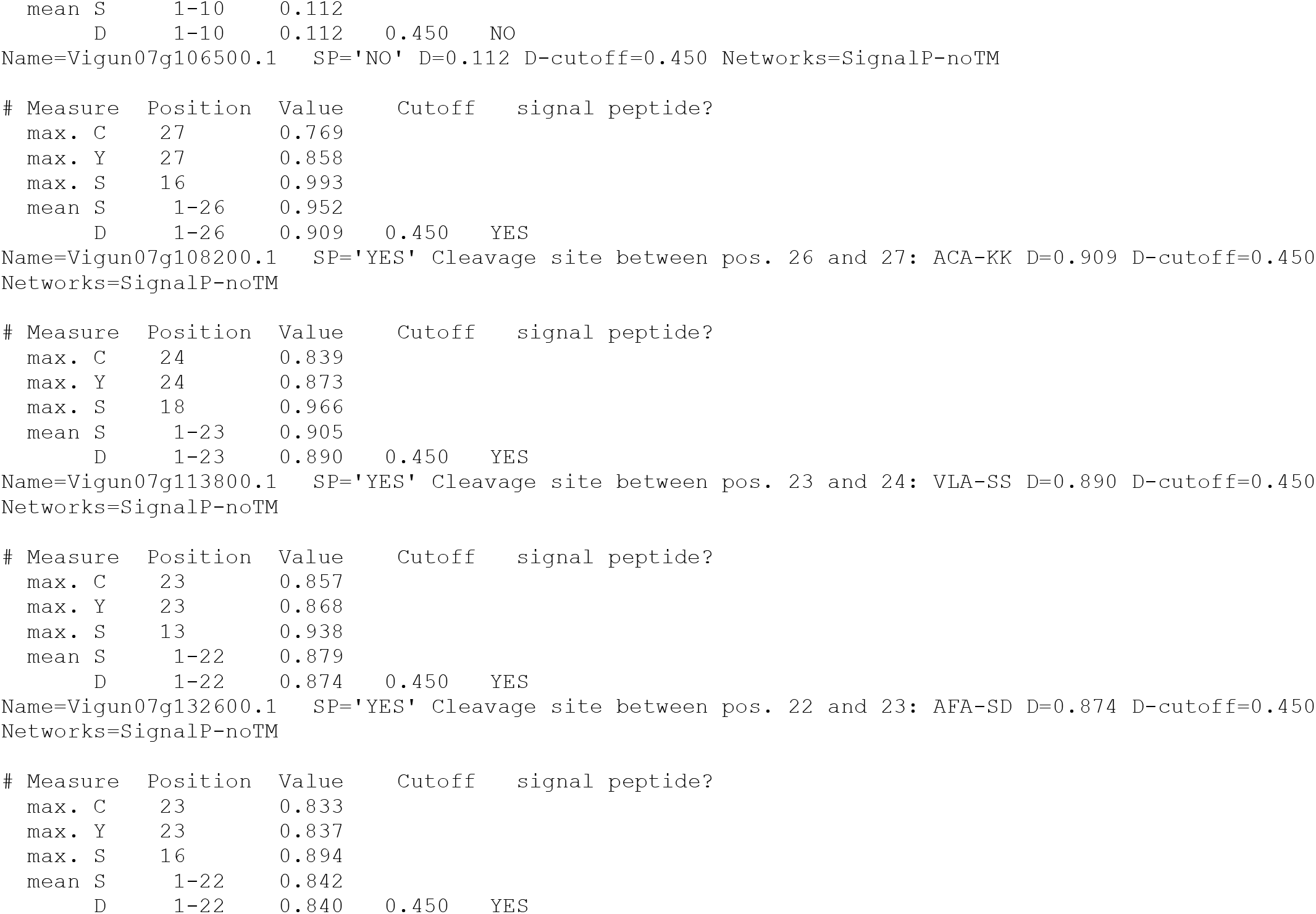
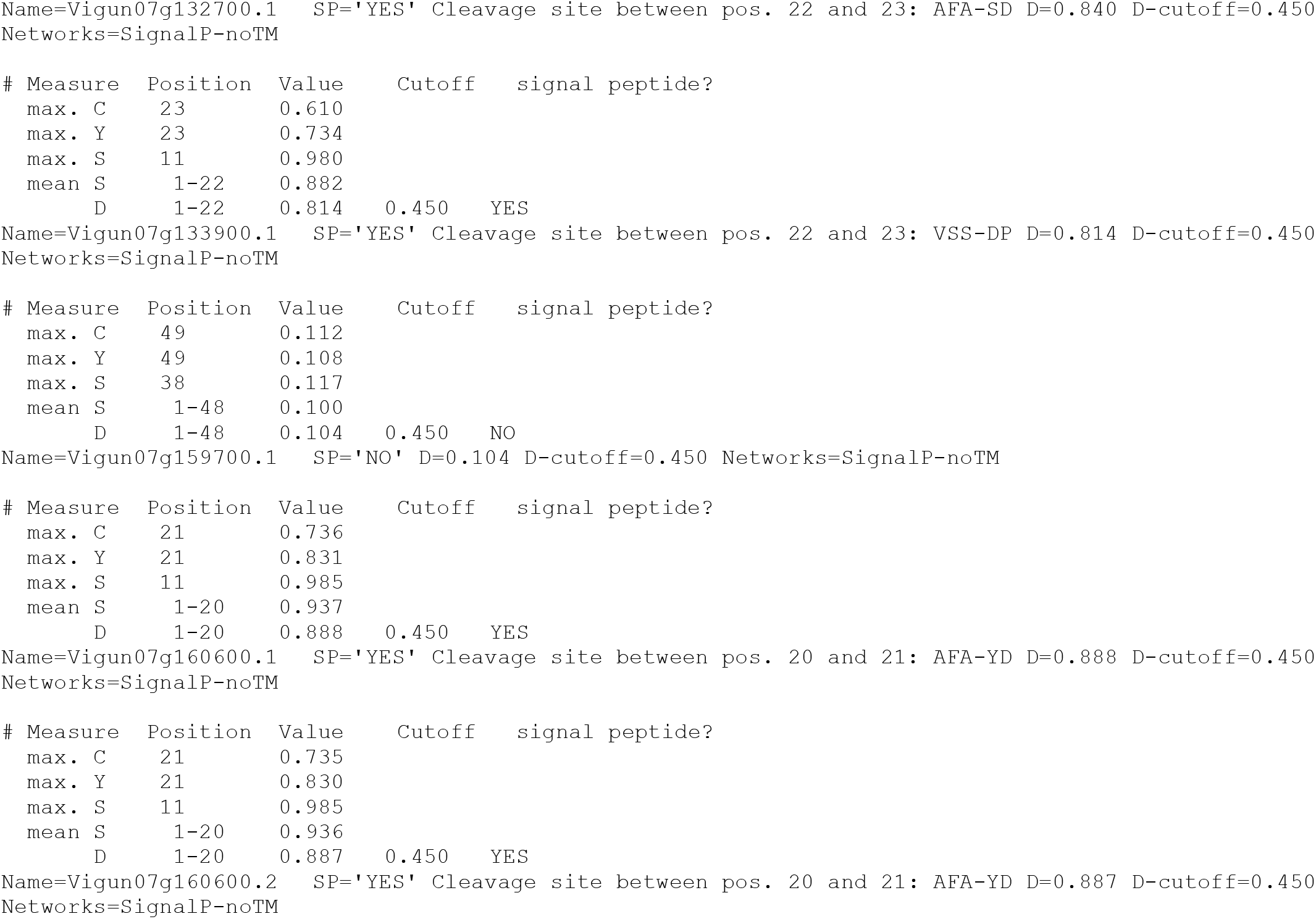
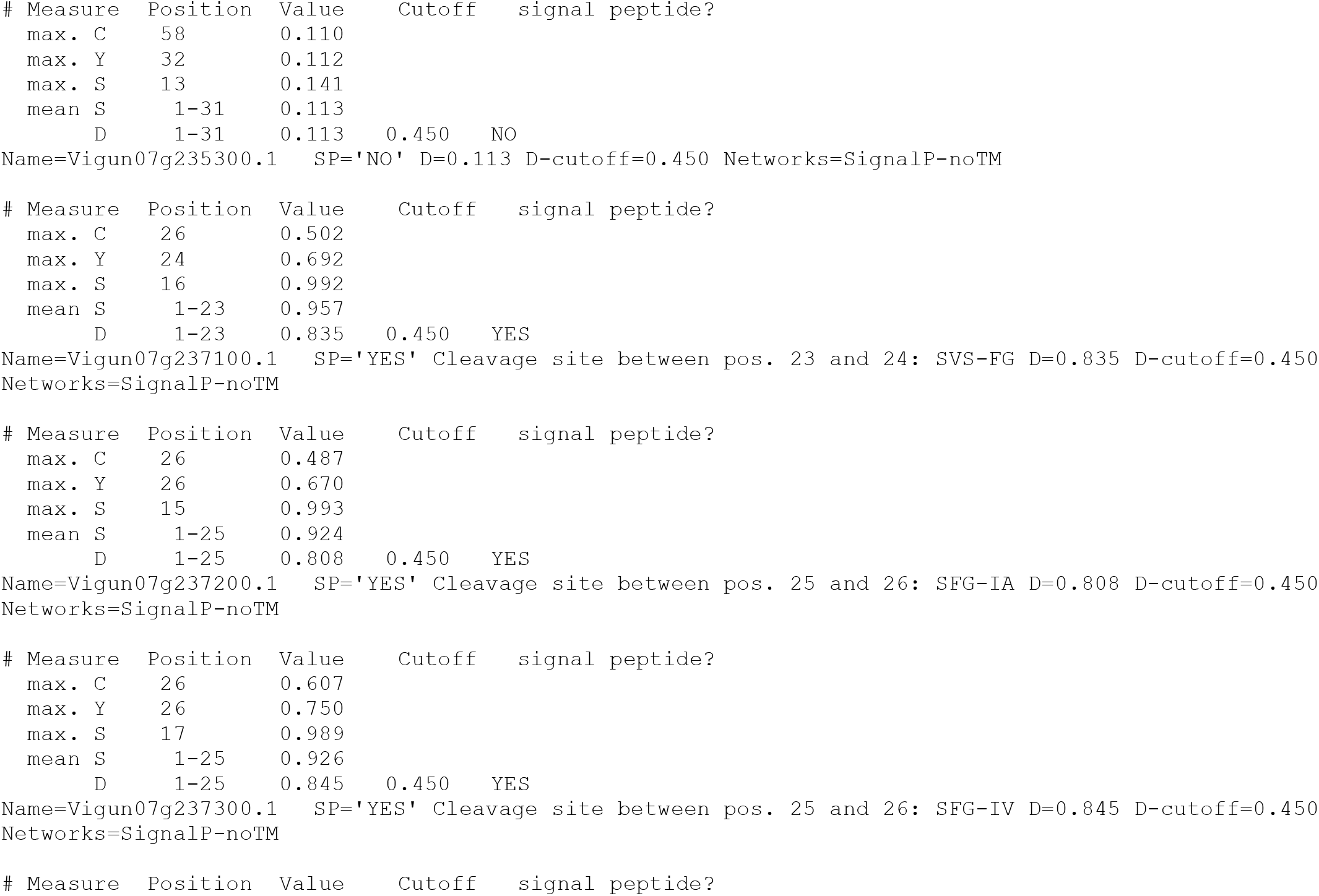
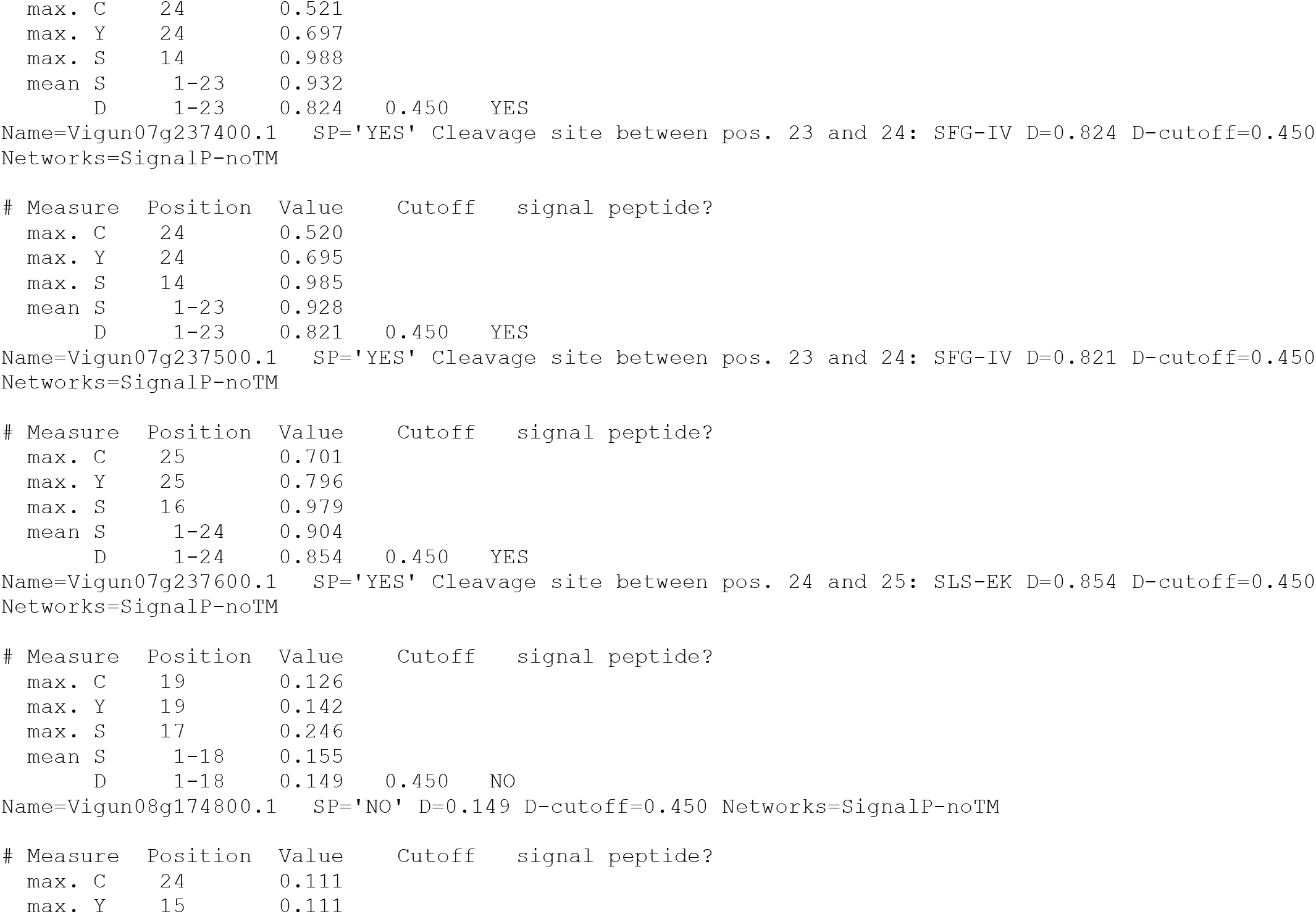
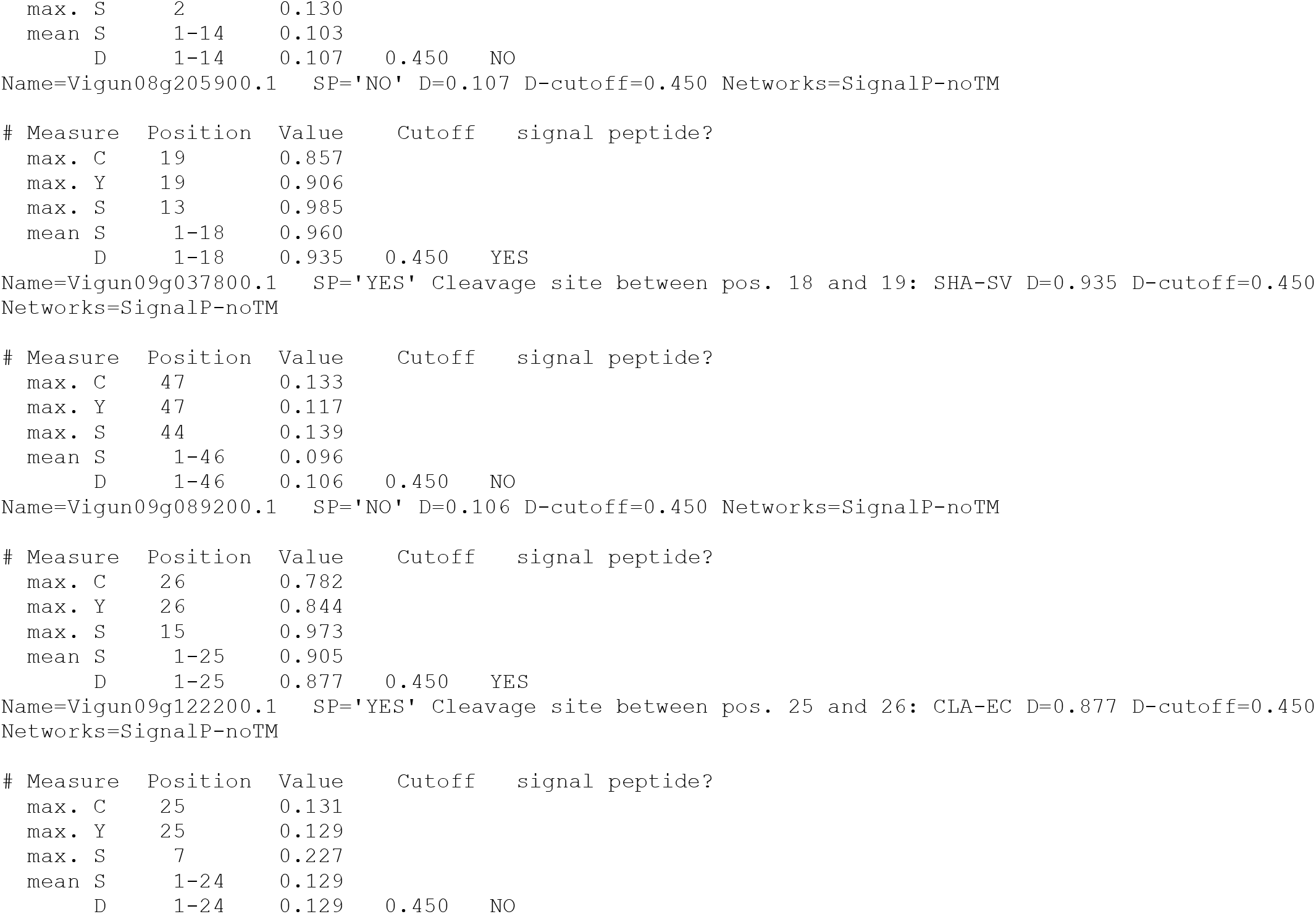
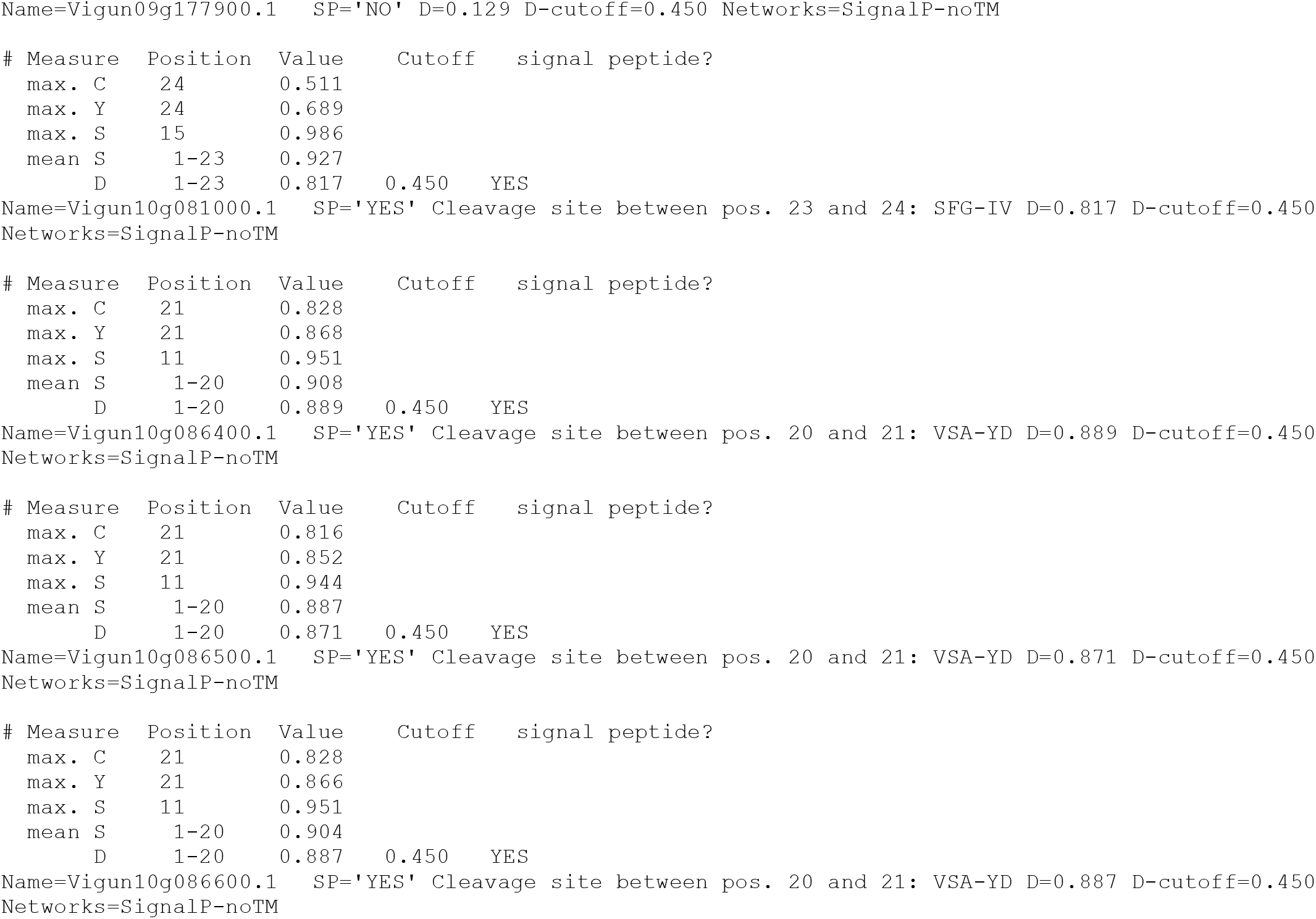
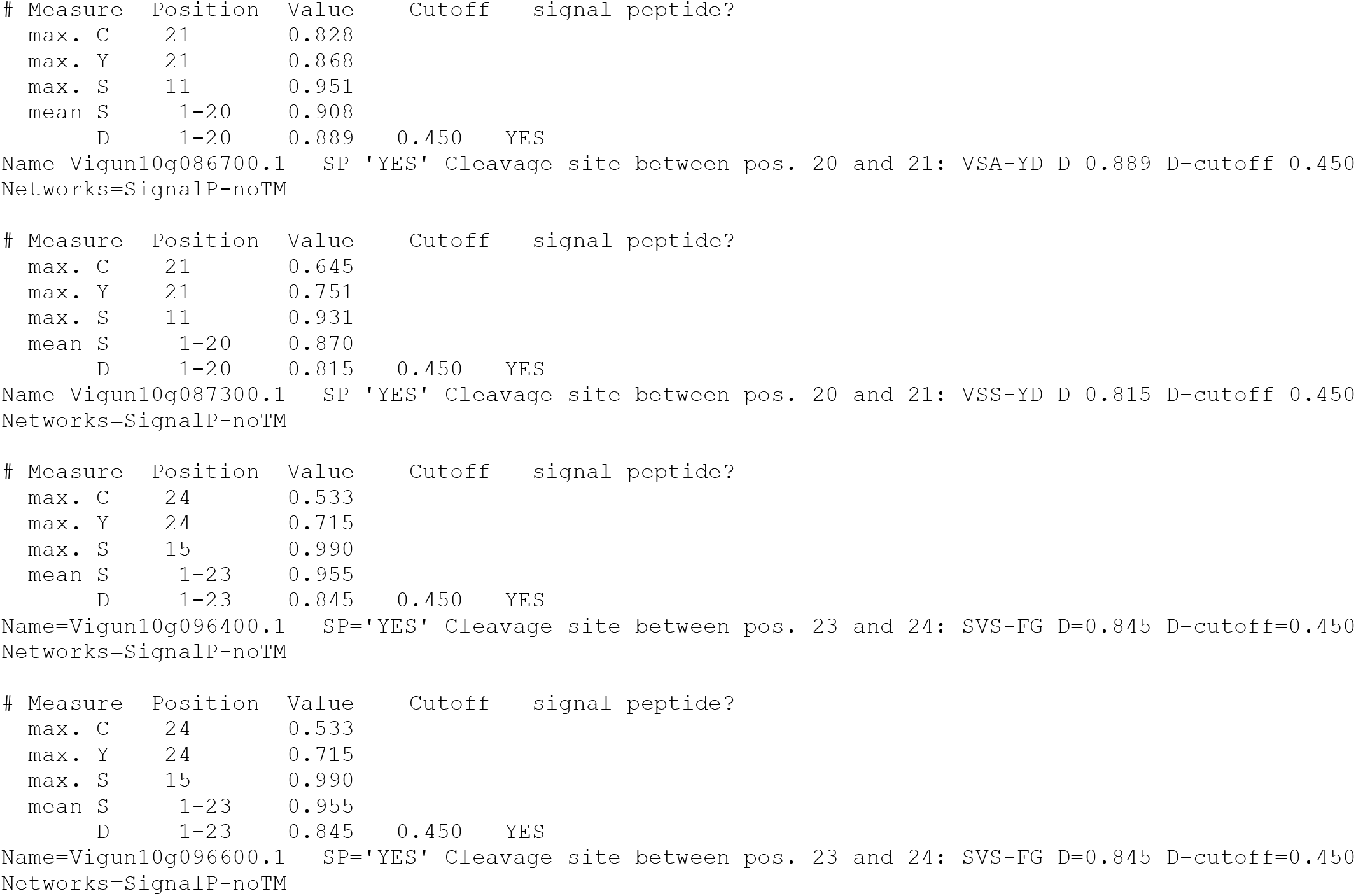
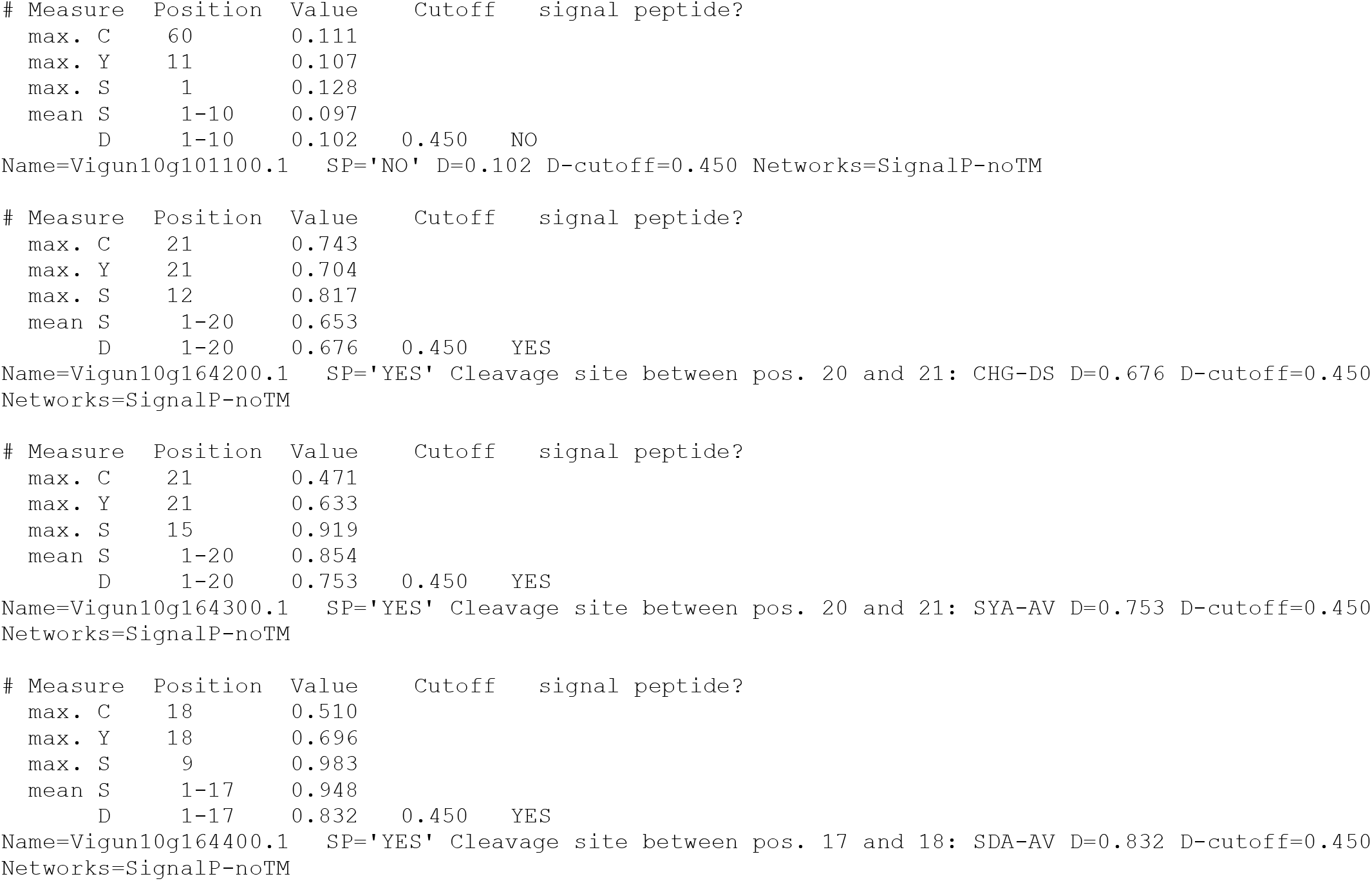
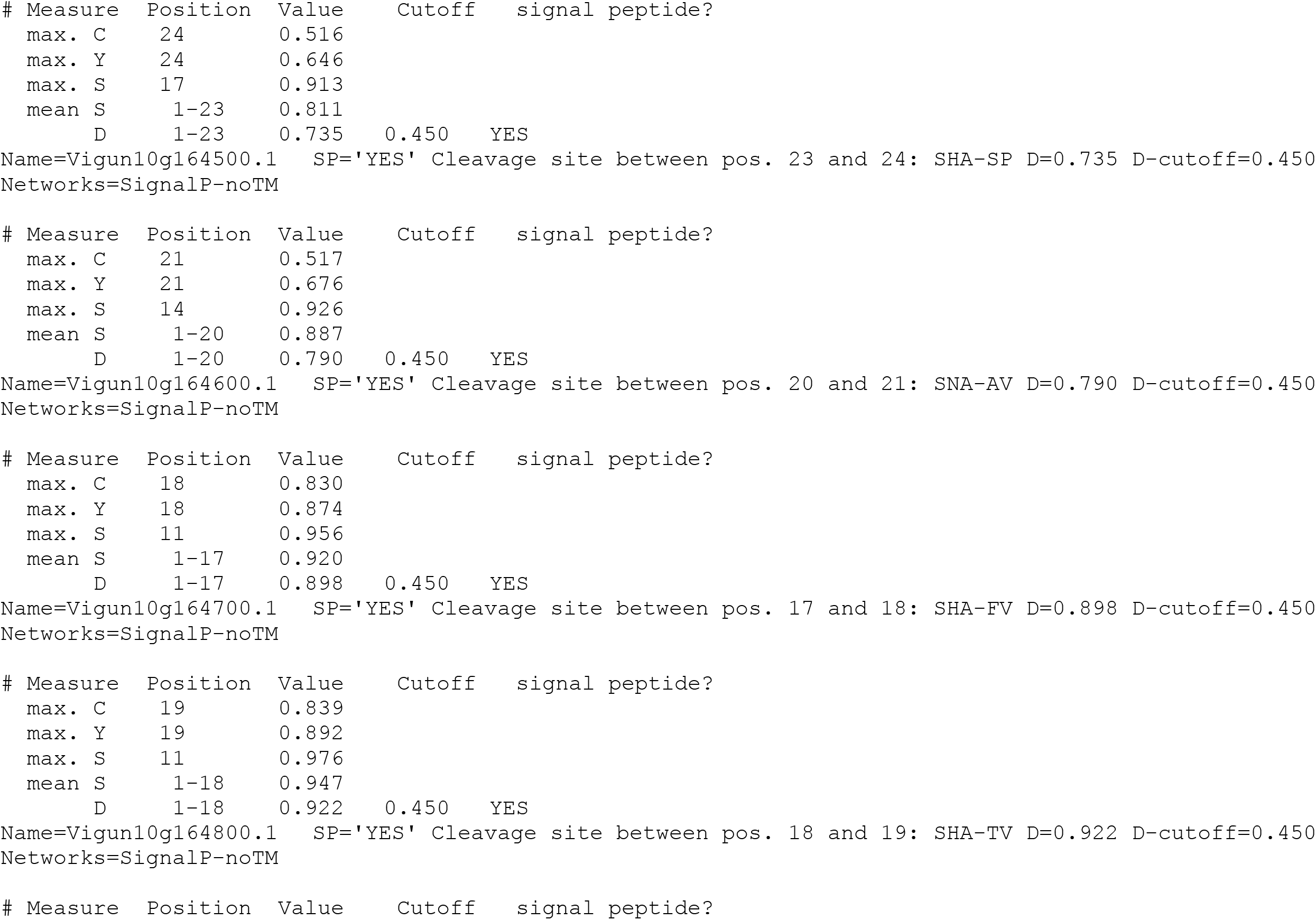
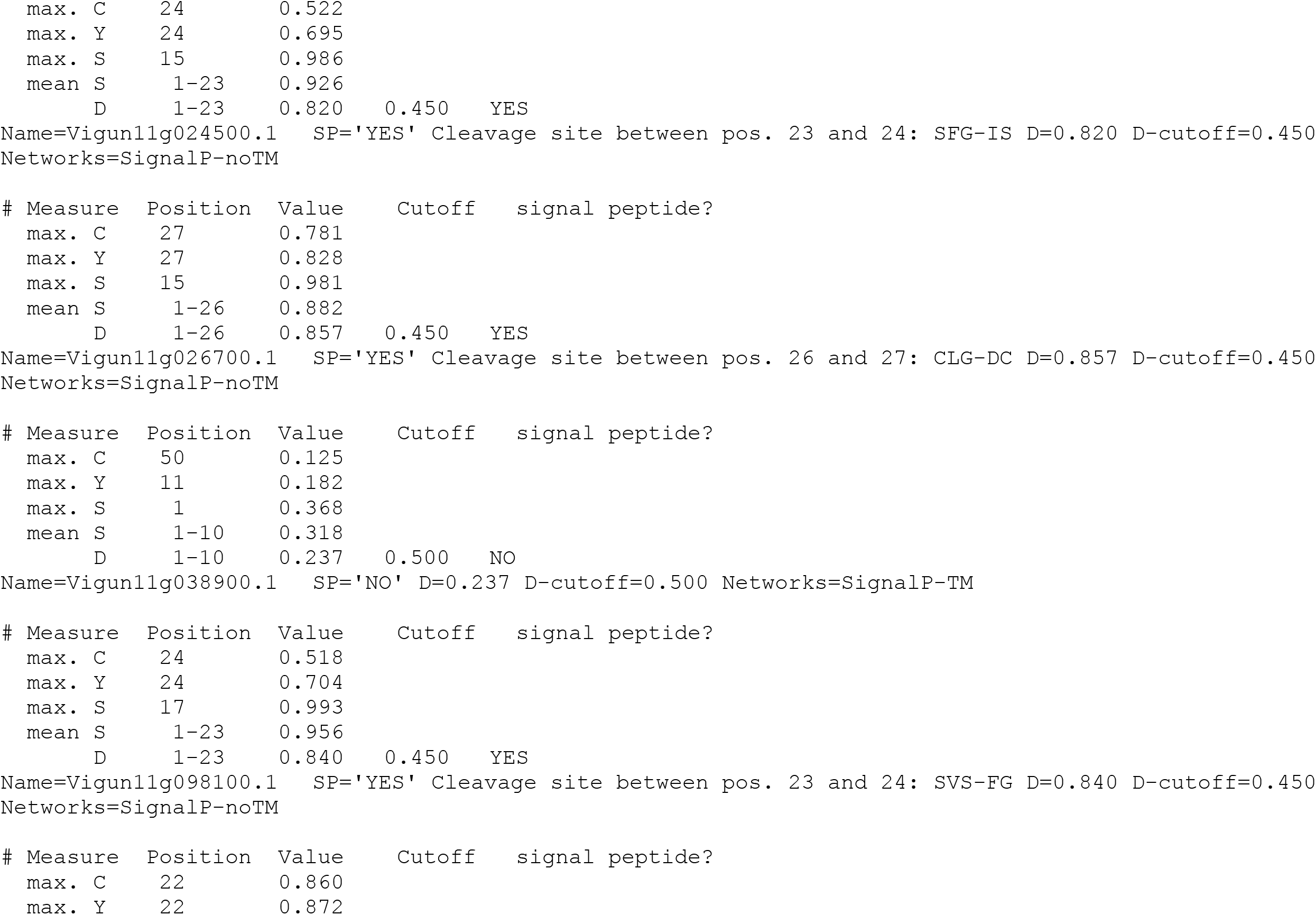
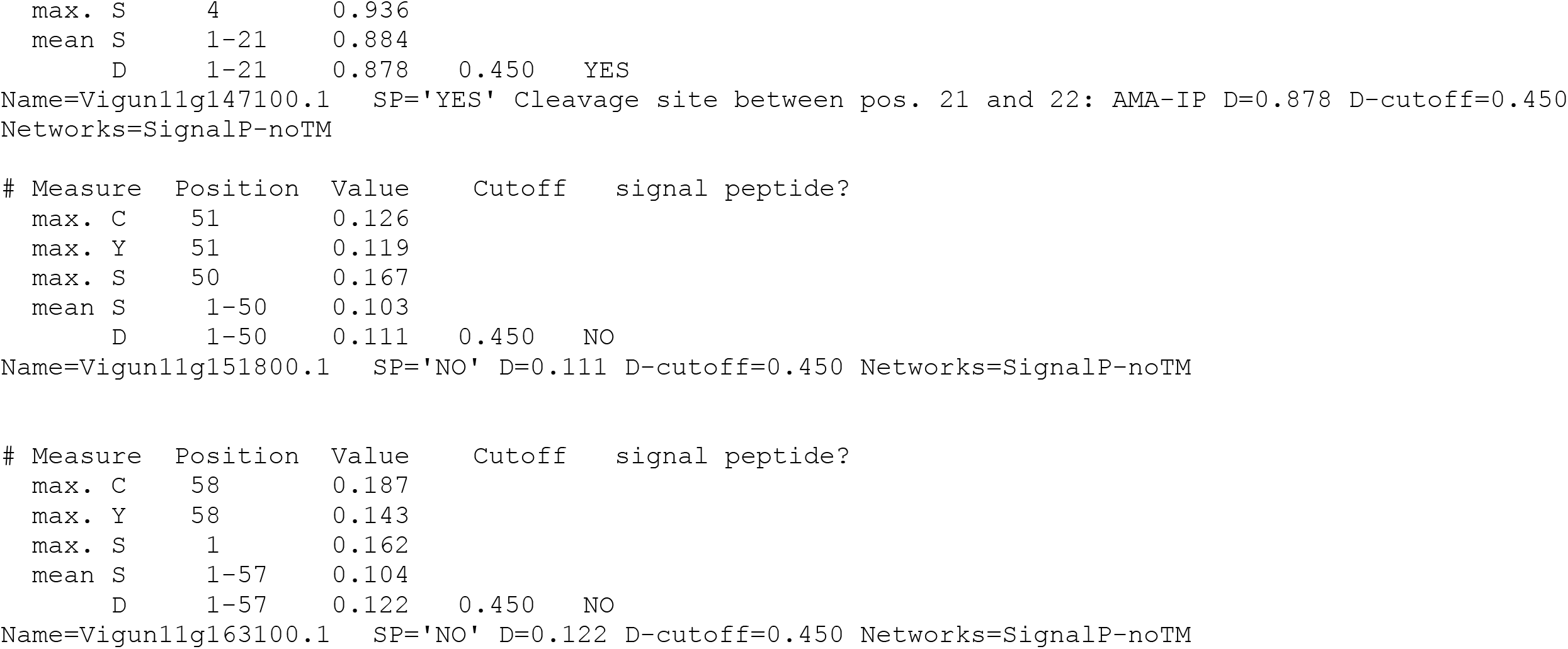
the 77 cupins sequences of the *V. unguiculata* showed the absence or presence of signal peptides and positions of cleavage sites.

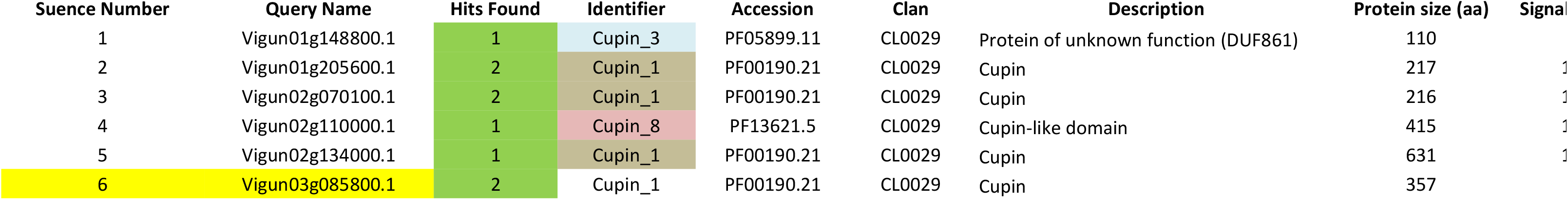
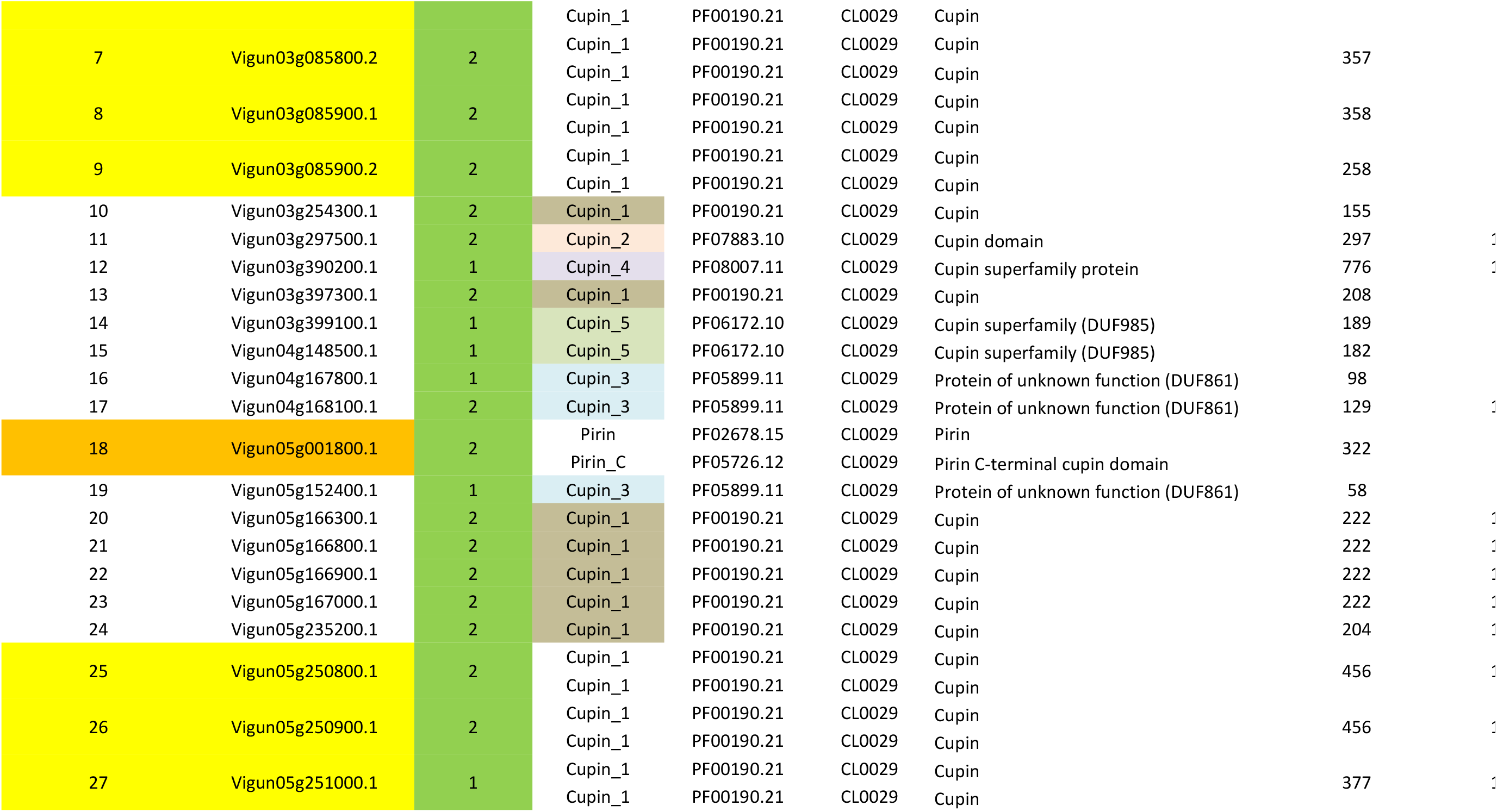
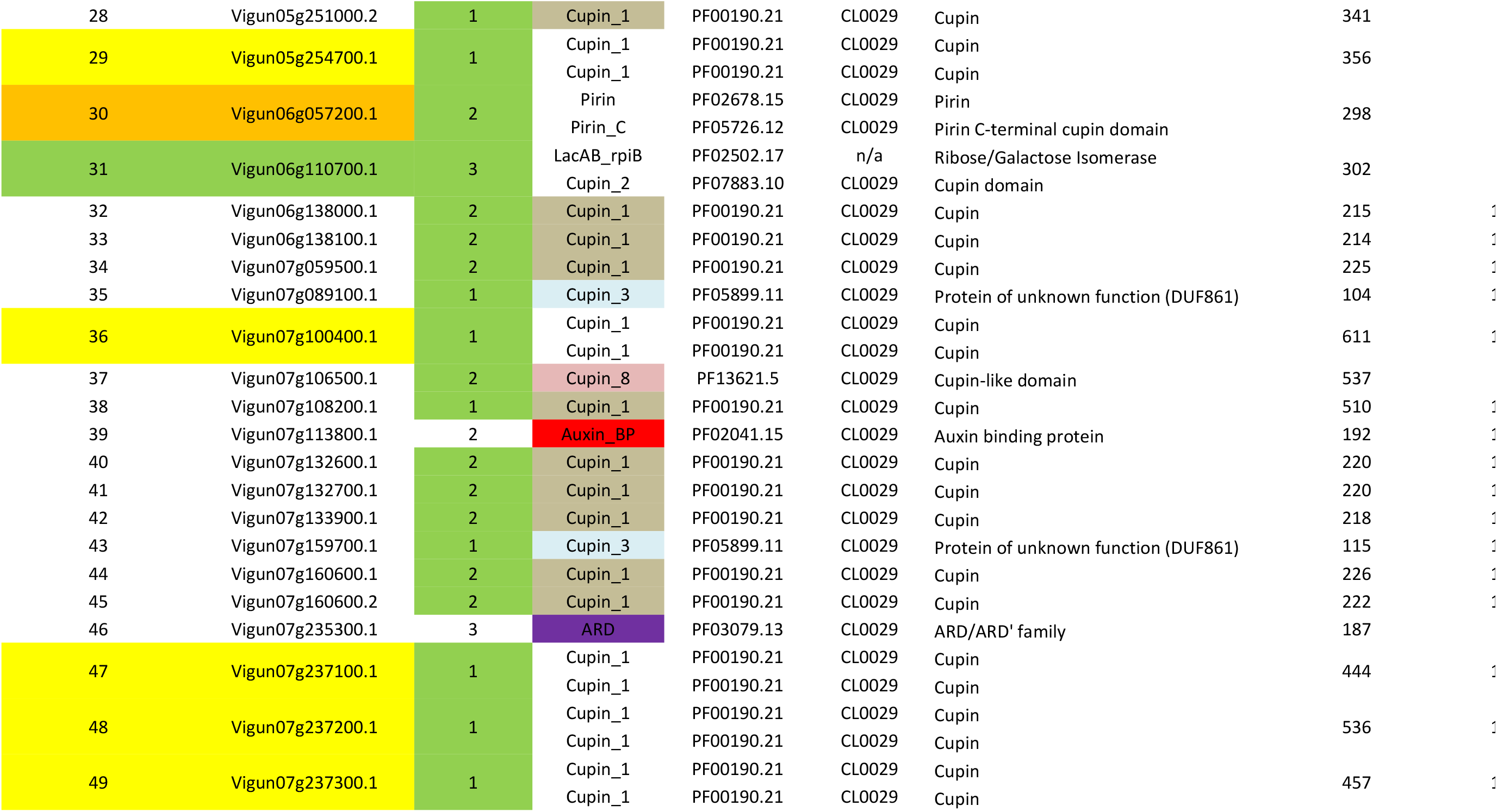
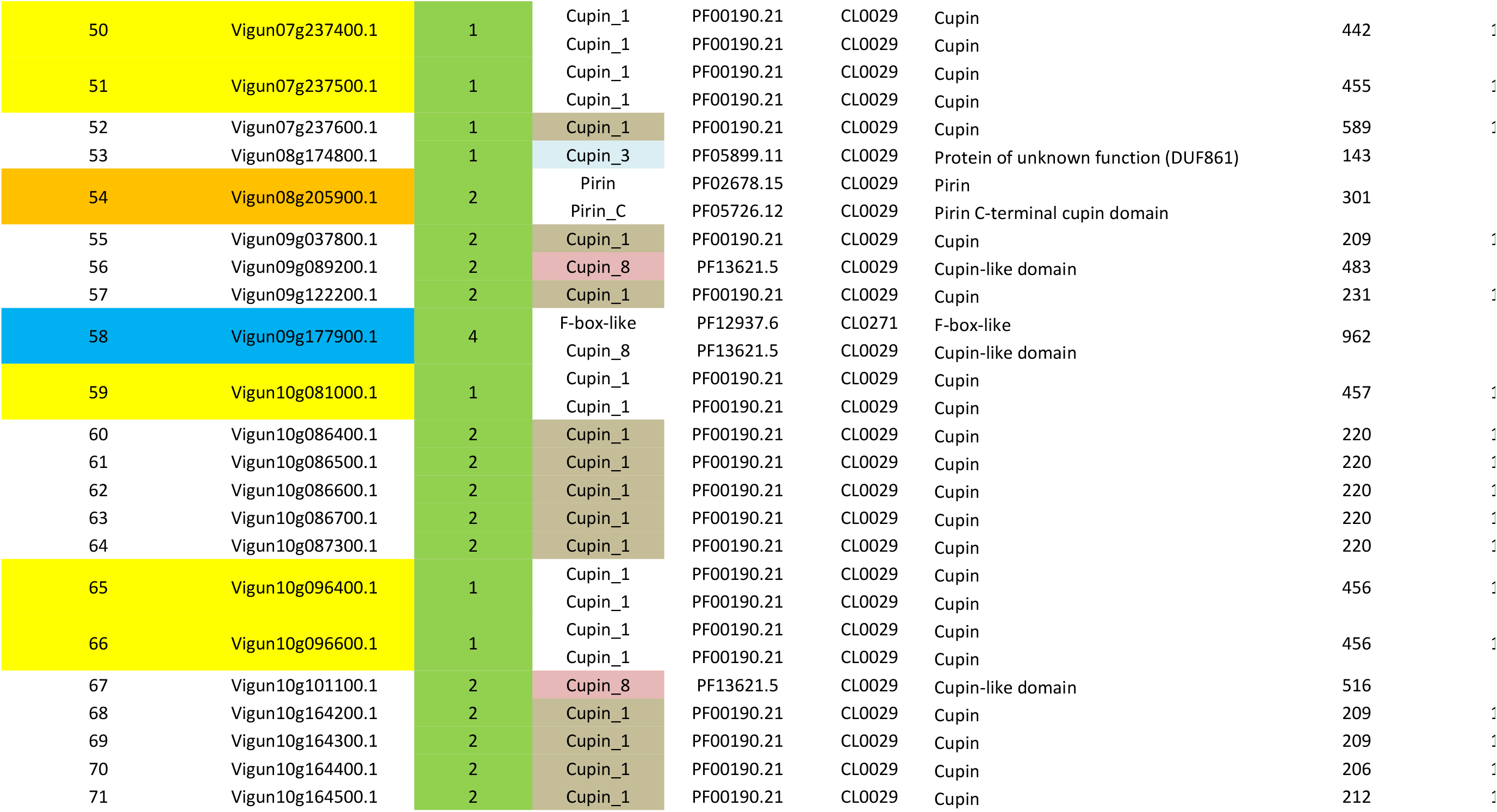
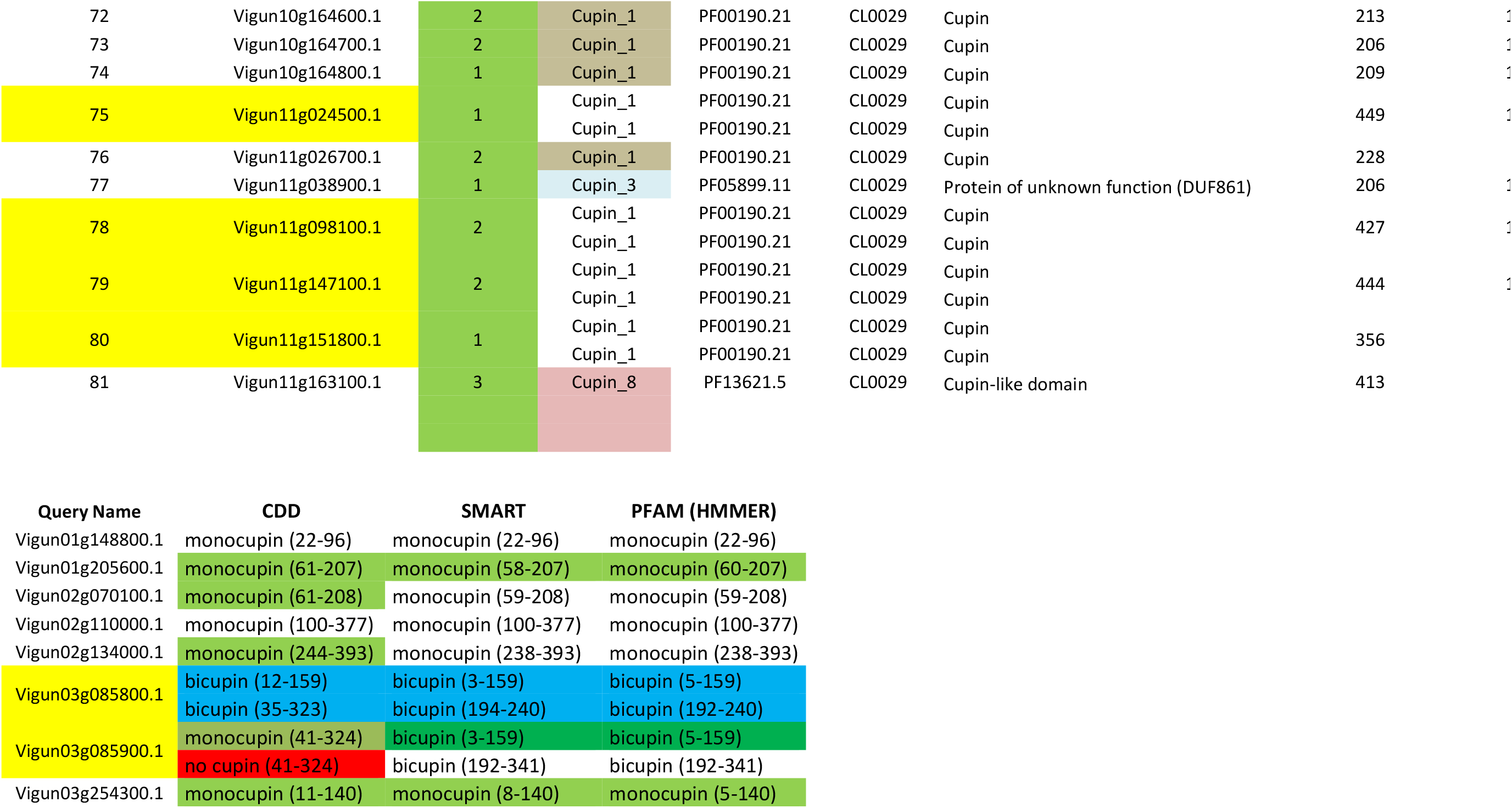
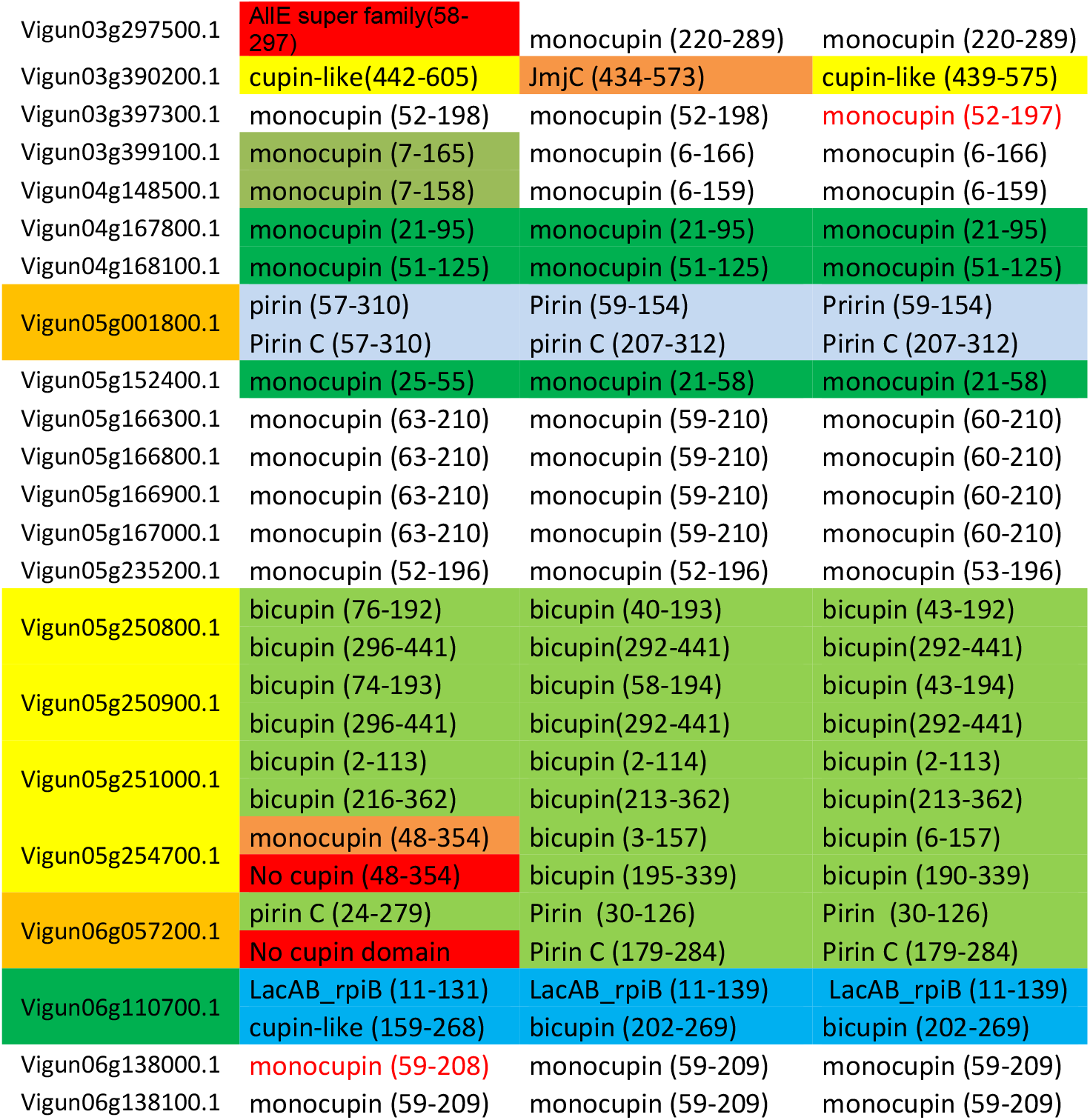
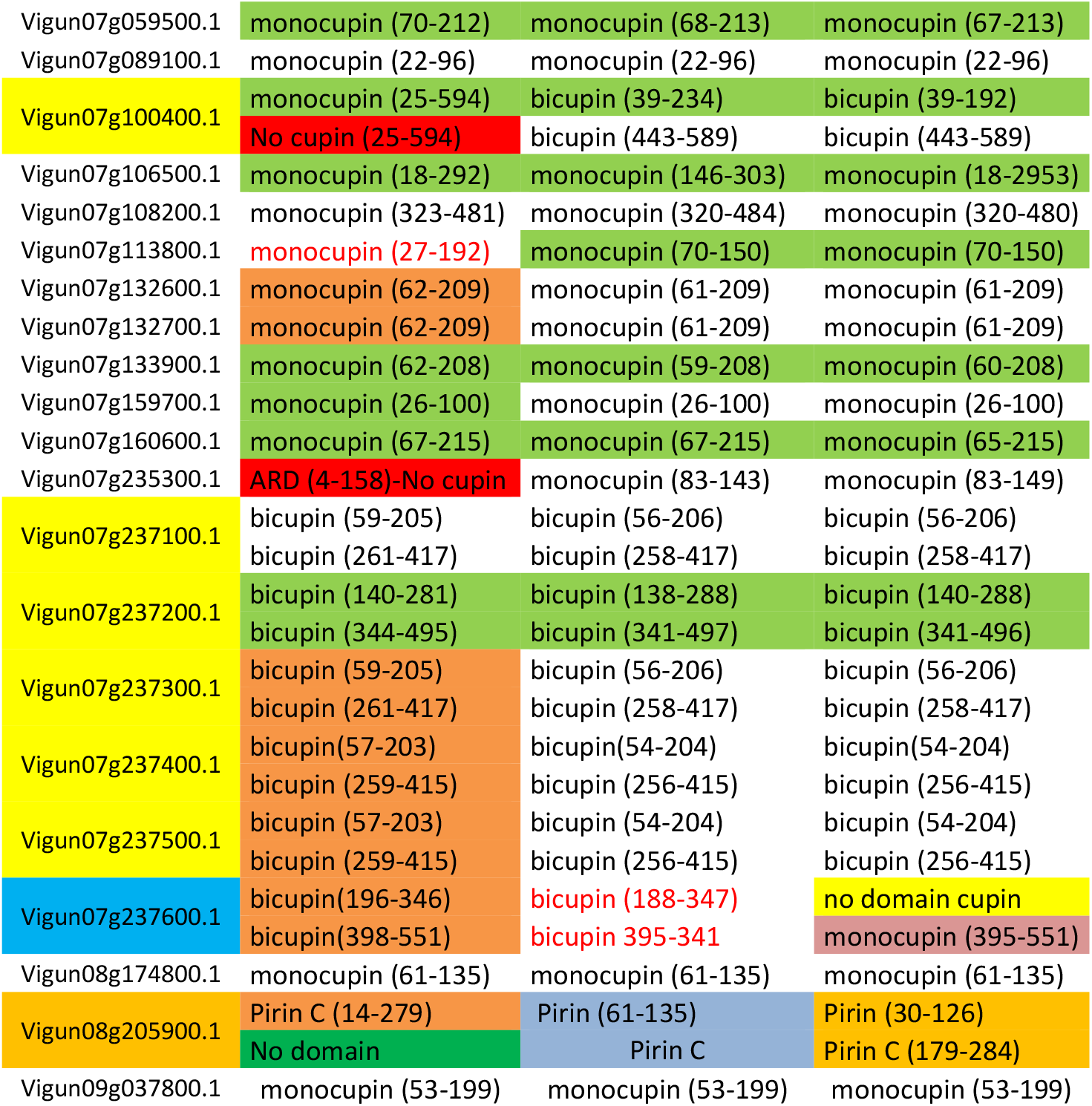
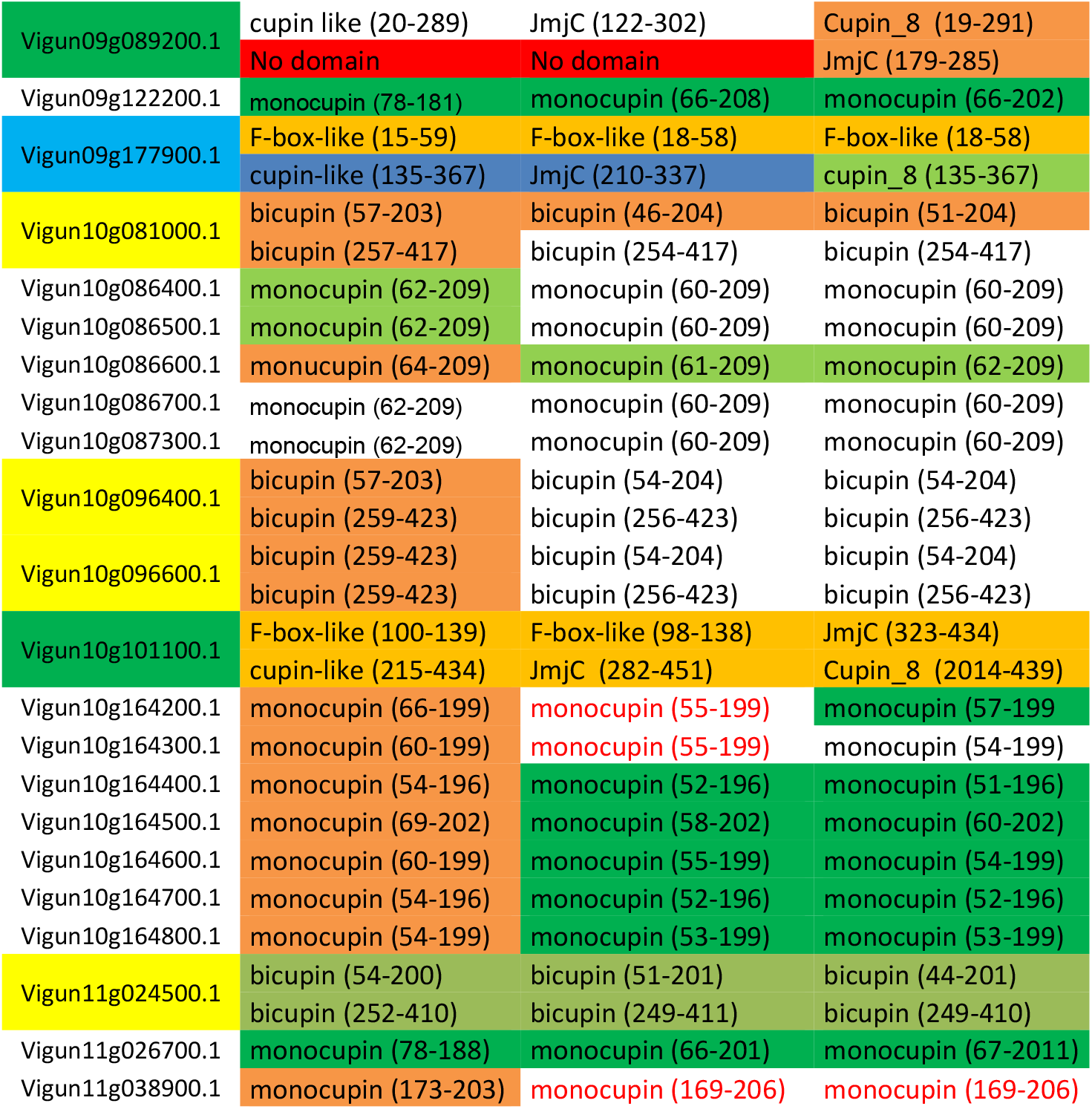
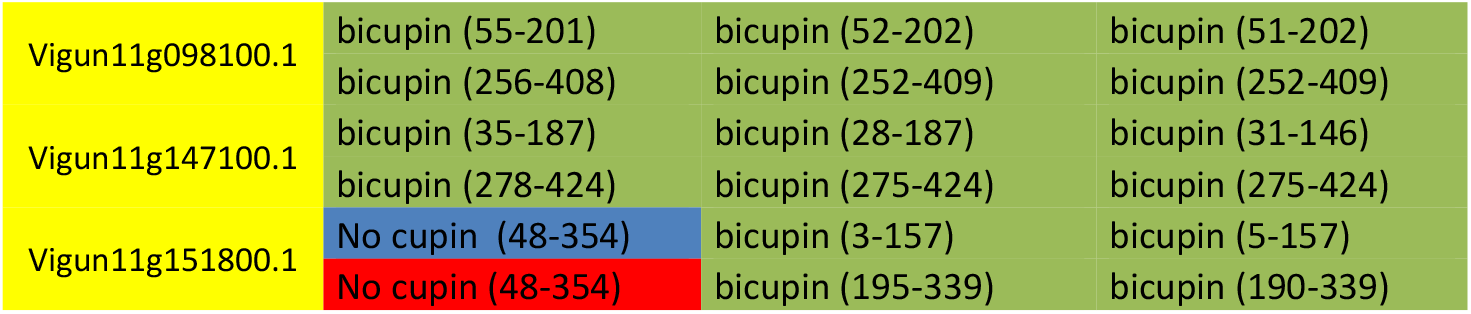

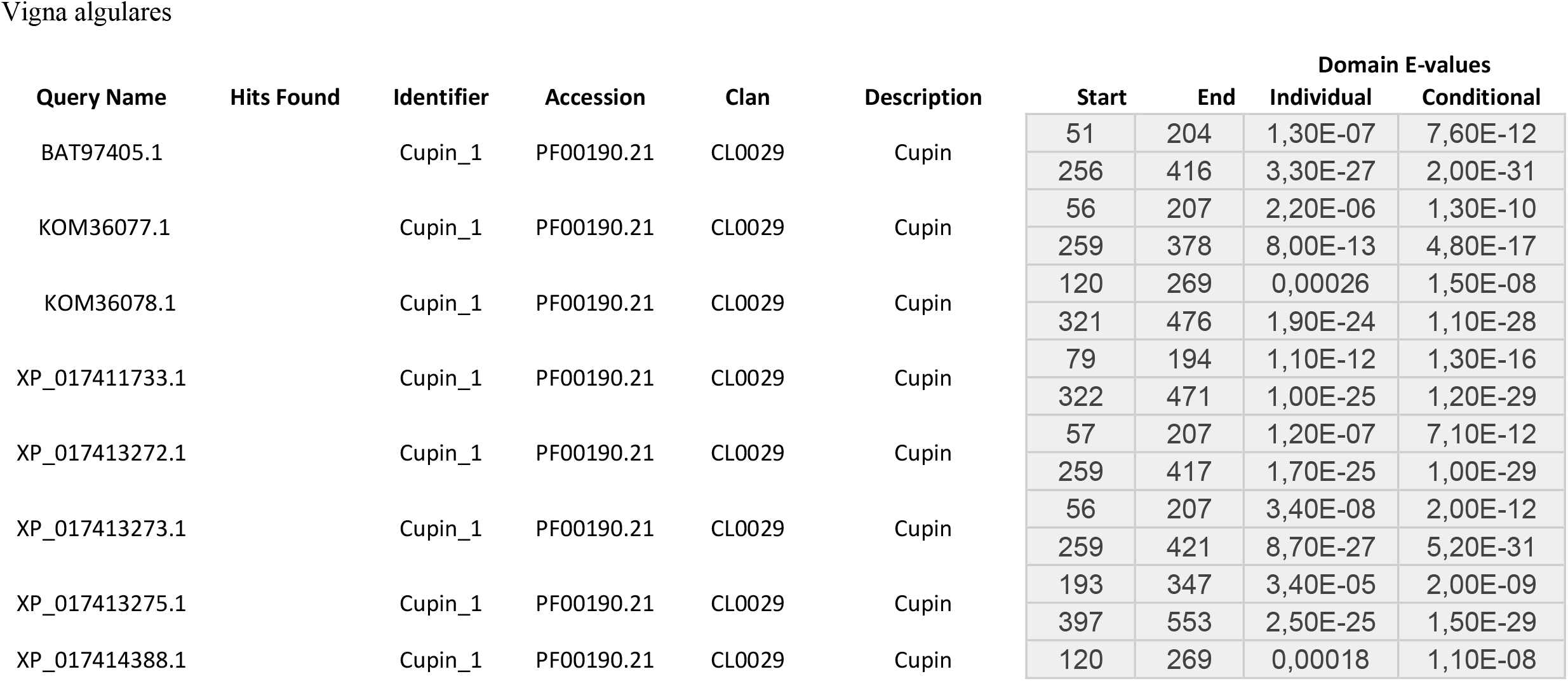
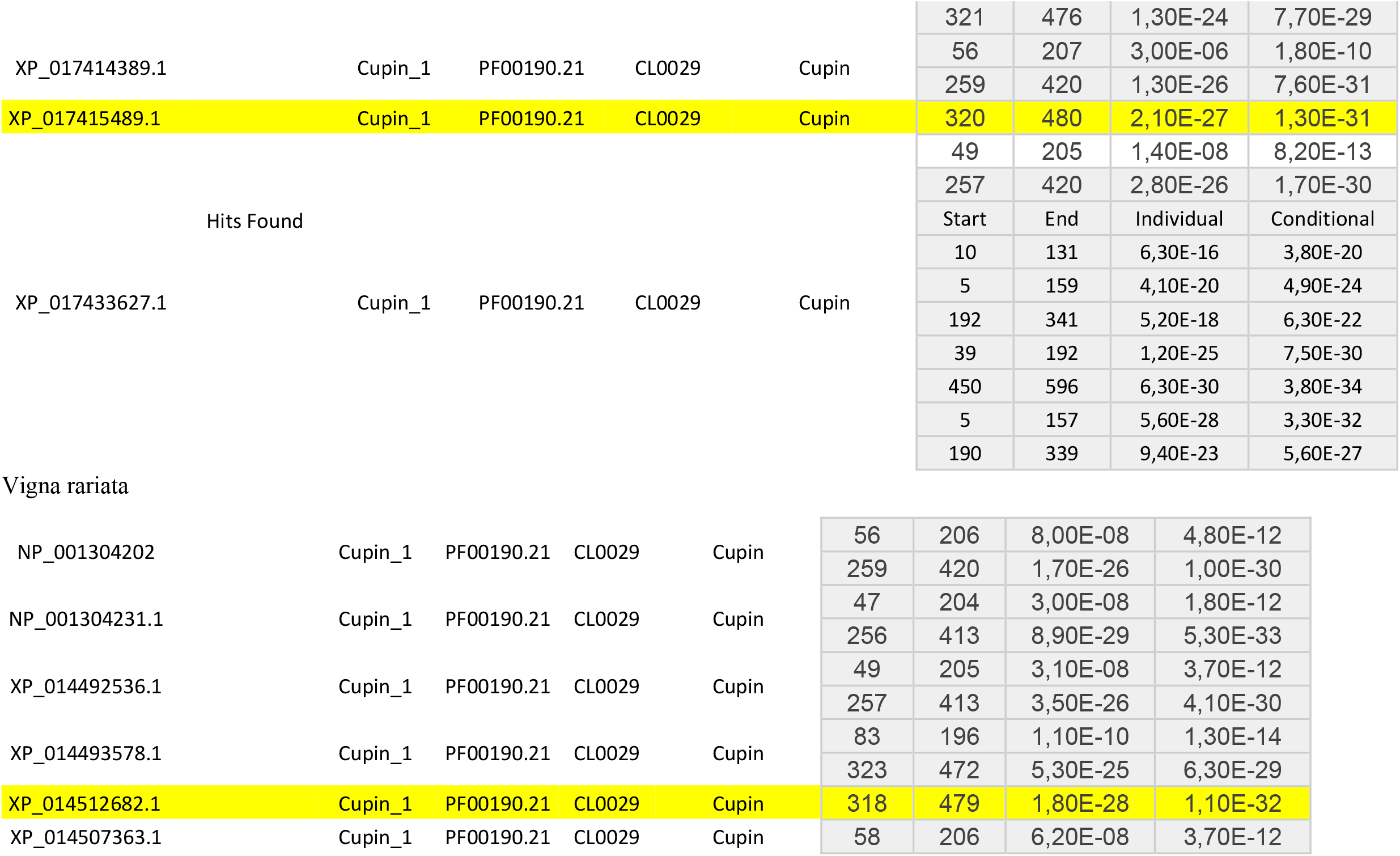
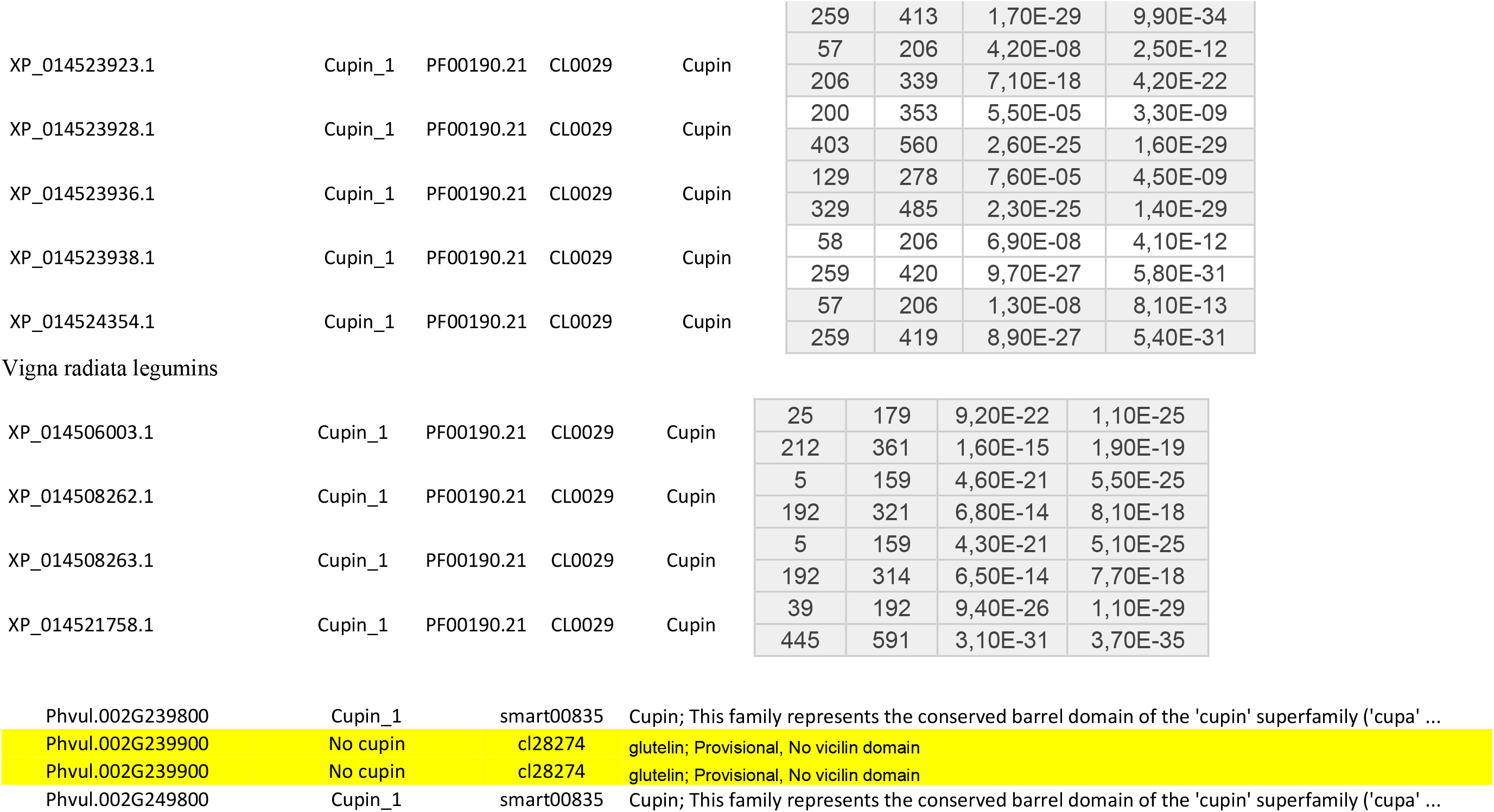
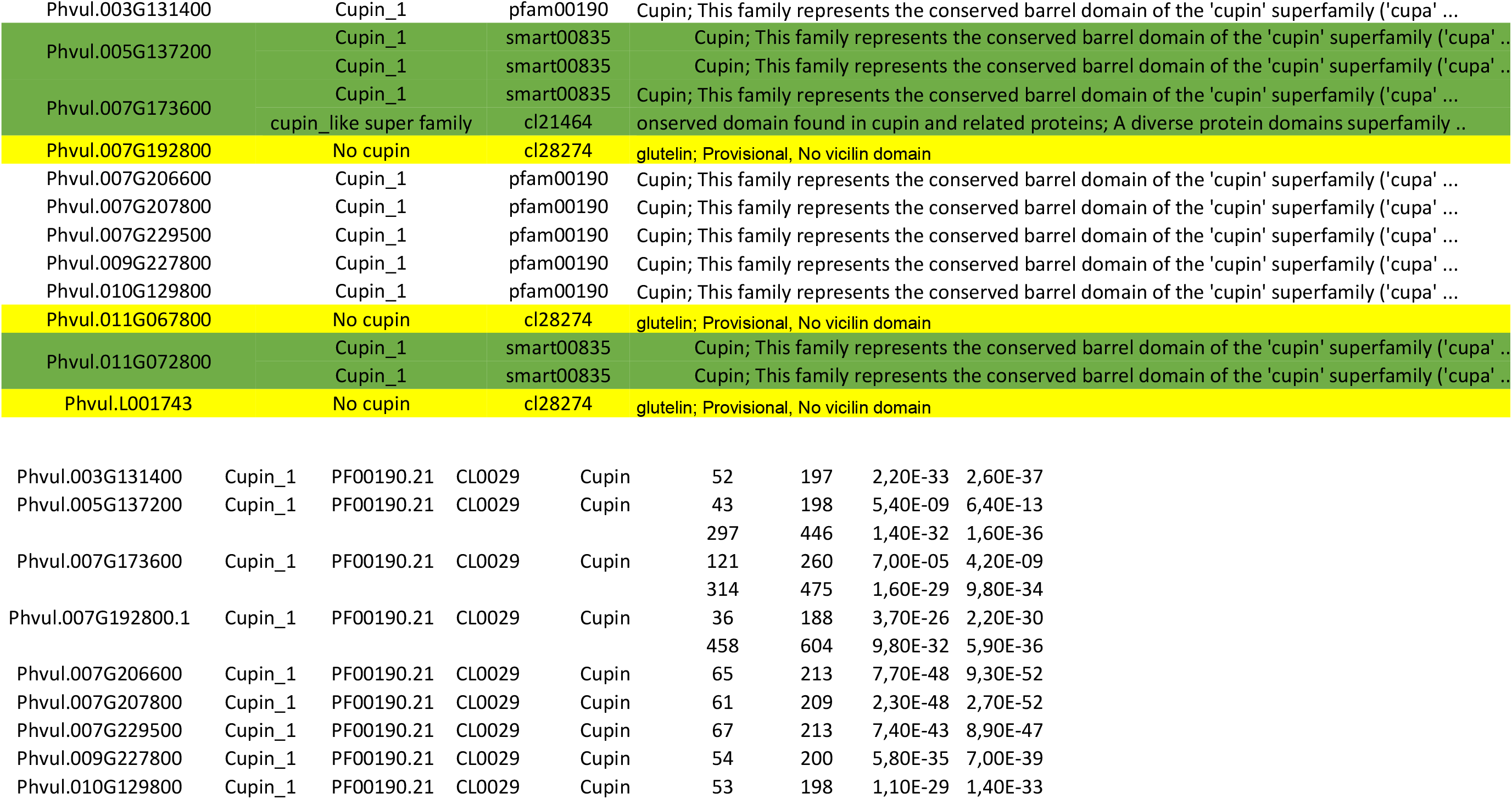
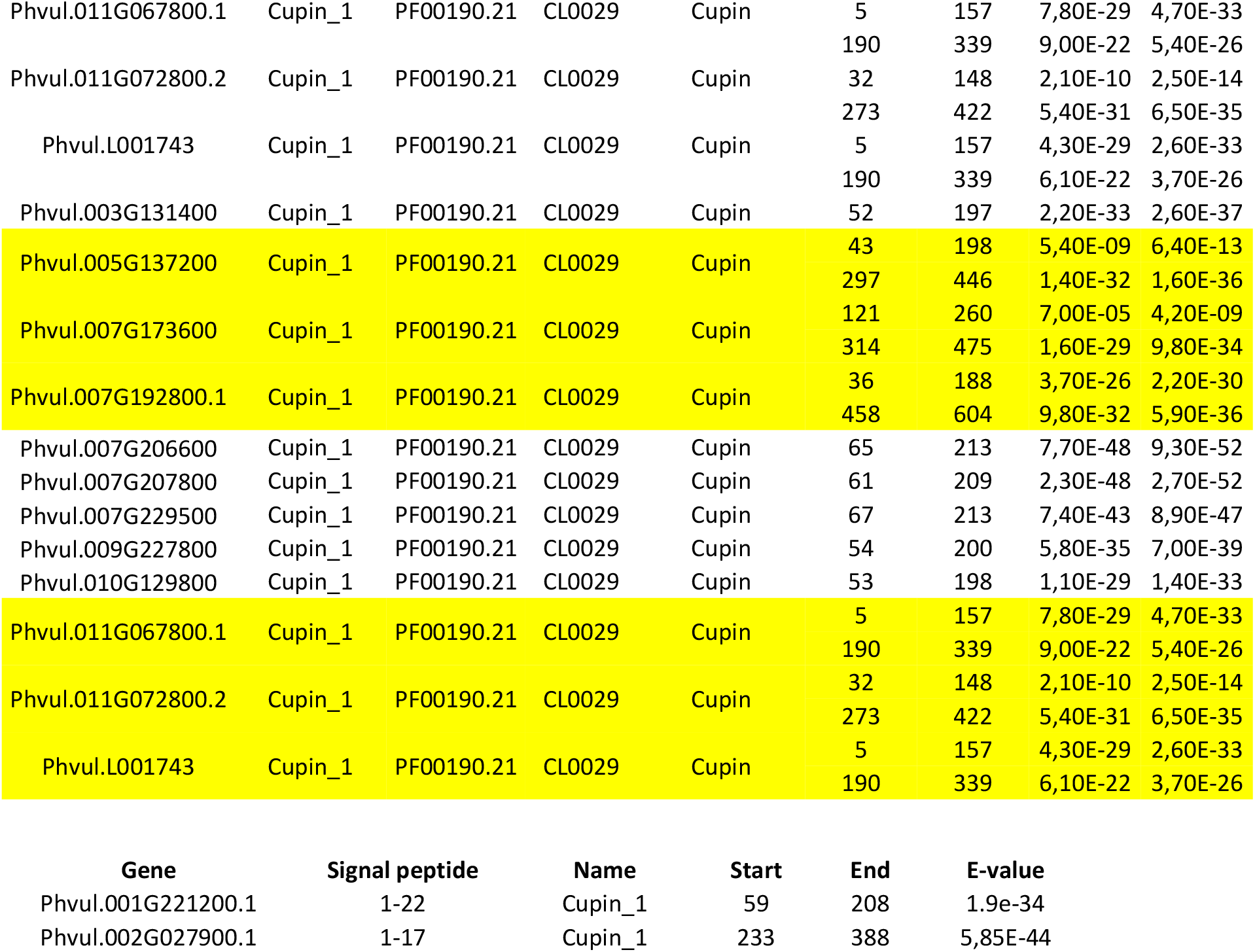
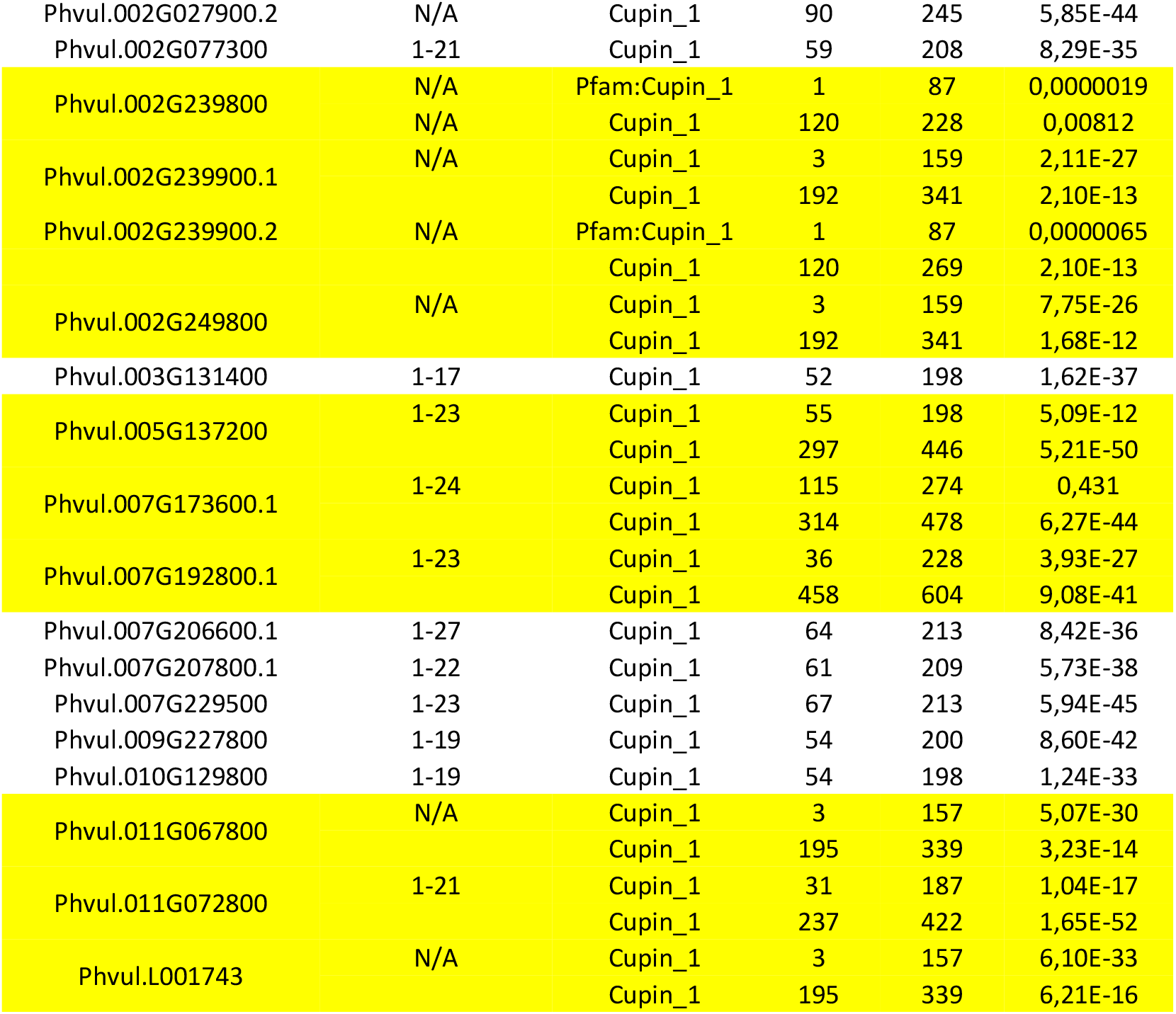

## References

Beilinson, V., Chen, Z., Shoemaker, R.C., Fischer, R.L., Goldberg, R.B., Nielsen, N.C.: Genomic organization of glycinin genes in soybean. - Theor. appl. Genet. 104: 1132–1140, 2002.

Chua, A.C.N., Hsiao, E.S.L., Yang, Y.C., Lin, L.J., Chou, W.M., Tzen, J.T.C.: Gene families encoding 11S globulin and 2S albumin isoforms of jelly fig (Ficus awkeotsang) achenes. - Biosci. Biotechnol. Biochem. 72: 506–513, 2008.

Domoney, C., Casey, R.: Measurement of gene number for seed storage proteins in Pisum. - Nucl. Acids Res. 13: 687–699, 1985.

Domoney, C., Ellis, T.H.N., Davies, D.R.: Organization and mapping of legumin genes in Pisum. - Mol. gen. Genet. 202: 280–285, 1986

Docimo, T., Caruso, I., Ponzoni, E., Mattana, M., Galasso, I., 2014. Molecular characterization of edestin gene family in Cannabis sativa. - Plant. Physiol. Biochem. 84: 142–148.

Dunwell, J.M., 1998. Cupins: a new superfamily of functionallydiverse proteins that include germins and plant seed storage proteins. Biotechnol. Genet. Engin. Rev. 15, 1–32.

Dunwell, J.M., Culham, A., Carter, C.E., Sosa-Aguirre, C.R., Goodenough, P.W., 2001. Evolution of functional diversity in the cupin superfamily. Trends Biochem. Sci. 26, 740–745.

Dunwell, J.M., Khuri, S., Gane, P.J., 2000. Microbial relatives of the seed storage proteins of higher plants: conservation of structure, and diversification of function during evolution of the cupin superfamily. Microbiol. Mol. Biol. Rev. 64, 153–179

Dunwell, J.M., 2002. Future prospects for transgenic crops. Phytochem. Rev. 1, 1–12.

Dunwell, J.M., 2003. Structure, function and evolution of the legumin seed storage proteins. In: Steinböchel, A., Fahnestock, S.R. (Eds.) Biopolymers, Vol. 8. Polyamides and Complex Proteinaceous Materials II. Wiley-VCH, Weinheim, pp. 223–253.

Ehlers, J.D., Hall, A.E., 1997. Cowpea (*Vigna unguiculata* L.Walp.), Field Crop Res. 53 187–204, https://doi.org/10.1016/S0378-4290(97)00031-2.

Finn, R.D., et al. (2016), The Pfam protein families database: towards a more sustainable future: Nucleic Acids Res. 2016 Jan 4; 44(Database issue): D279–D285. https://doi.org/101093/nar/gkv1344

Goodstein, M.D., et al., 2012. Phytozome: a comparative platform for green plant genomics, Nucleic Acids Res. 2012 40 (D1): D1178–D1186.

Howe, R.W., Currie, J.E., 1964. Some laboratory observations on the rates of evelopment, mortality and oviposition of several species of Bruchidae breeding in stored pulses, Bull. Entomol. Res. 55 437, https://doi.org/10.1017/S0007485300049580

Marchler-Bauer, A., et al. 2017. “CDD/SPARCLE: functional classification of proteins via subfamily domain architectures, Nucleic Acids Res. 45(D) 200–3.

Osborne, N.J., et al. Prevalence of challenge-proven IgE-mediated food allergy using population-based sampling and predetermined challenge criteria in infants. J Allergy Clin Immunol 2011; 127:668–676.e1-2. https://doi.org/1016/j.jaci.2011.01.039

Pang, H., et al., 2014. Crystal Structure of Human Pirin. An iron-binding nuclear protein and transcription cofactor. 279: 1491–1498. https://doi.org/10.1074/jbc.M310022200.

Ponzoni, E., Brambilla, I.M., Galasso, I., 2018. Biol Plant 62: 693. https://doi.org/10.1007/s10535-018-0810-7

Potter, S.C., et al., 2018. Nucleic Acids Research. Web Server Issue 46: W200–W204.

Petersen, T.N., 2011. SignalP 4.0: discriminating signal peptides from transmembrane regions. Nat Methods. 2011 Sep 29;8(10):785–6. doi: 10.1038/nmeth.1701

Kriz, A.L., (1999) 7S Globulins of Cereals. In: Shewry P.R., Casey R. (eds) Seed Proteins. Springer, Dordrecht. https://doi.org/10.1007/978-94-011-4431-5_20

Singh, B.B., Ajeigbe, H.A., Tarawali, S.A., Fernandez-Rivera, S., Abubakar, M., 2003. Improving the production and utilization of cowpea as food and fodder, Field Crop Res. 84 169–177, https://doi.org/10.1016/S0378-4290(03)00148-5

Rocha, A.J., 2018. Cloning of cDNA sequences encoding cowpea (Vigna unguiculata) vicilins: Computational simulations suggest a binding mode of cowpea vicilins to chitin oligomers, International Journal of Biological Macromolecules, 117: 565–573 https://doi.org/10.1016/j.ijbiomac.2018.05.197.

Sales, M.P., Macedo, M.R.L., Xavier-Filho, J, 1992. Digestibility of cowpea (Vigna unguiculata) vicilins by pepsin, papain and bruchid (insect) midgut proteinases, Comp. Biochem. Physiol. B Biochem. Mol. Biol. 103: 945–950, https://doi.org/10.1016/0305-0491(92)90220-L.

Sales, M.P., Pimenta, P.P., Paes, N.S., Grossi-De-Sa, M.F., Xavier J., 2001. Vicilins (7S storage globulins) of cowpea (Vigna unguiculata) seeds bind to chitinous structures of the midgut of Callosobruchus maculatus (Coleoptera: Bruchidae) larvae, Braz. J. Med. Biol. Res. 34: 27–34, https://doi.org/10.1590/S0100-879X2001000100003.

Saitou, N., Nei M., 1987. The neighbor-joining method: a new method for reconstructing phylogenetic trees. Molecular Biology and Evolution, 4:.406–425.

Shotwell, M.A., Larkins, B.A., 2012. Improvement of the protein quality of seeds by genetic engineering. - In: Dennis, E.S., Lewellyn, D.J. (ed.): Molecular Approaches to Crop Improvement. Pp. 33–61. Springer-Verlag, Wien - New York 2012.

Schultz, J., et al., 2000. SMART: A Web-based tool for the study of genetically mobile domains. Nucleic Acids Res. 28: 231–234.

Sreedhar, R., Kaul, P., 2016. Cupincin: A Unique Protease Purified from Rice (Oryza sativa L.) Bran Is a New Member of the Cupin Superfamily. PLoS ONE 11(4): e0152819. doi:10.1371/journal.pone.0152819.

Tamakura, K. et al., 2016. MEGA7: molecular evolutionary genetics analysis using maximum likelihood, evolutionary distance, and maximum parsimony methods. Molecular Biology and Evolution 33 (7):1870–4. https://doi.org/10..1093/molbev/msw054

